# Cell-based optimisation and characterisation of genetically encoded, location-based biosensors for Cdc42 or Rac activity

**DOI:** 10.1101/2022.09.12.507573

**Authors:** Eike K. Mahlandt, Gabriel Kreider-Letterman, Anna O. Chertkova, Rafael Garcia-Mata, Joachim Goedhart

**Author notes:** Correspondence: | Twitter: @joachimgoedhart | ORCID: 0000-0002-0630-3825.

## Abstract

Rac and Cdc42 are Rho GTPases which regulate the formation of lamellipoda and filopodia and are therefore crucial in processes such as cell migration. Relocation-based biosensors for Rac and Cdc42 have not been characterized well in terms of their specificity or affinity. In this study, we identify relocation sensor candidates for either Rac or Cdc42. We compared their (i) ability to bind the constitutively active Rho GTPases, (ii) specificity for Rac and Cdc42 and (iii) relocation efficiency in cell-based assays. Subsequently, the relocation efficiency was improved by a multi-domain approach. For Rac1 we found a sensor candidate with low relocation efficiency. For Cdc42 we found several sensors with sufficient relocation efficiency and specificity. These optimized sensors enable the wider application of Rho GTPase relocation sensors, which was showcased by the detection of local endogenous Cdc42 activity at assembling invadopodia. Moreover, we tested several fluorescent proteins and HaloTag for their influence on the recruitment efficiency of the Rho location sensor, to find optimal conditions for a multiplexing experiment. The characterization and optimization of relocation sensors will broaden their application and acceptance.

## Introduction

The morphology of the cell is mainly defined by the cytoskeleton. The cytoskeleton in mammalian cells consists of a network of different protein polymers and one of those is the actin cytoskeleton. This actin network is dynamic and constantly assembling and disassembling, adjusting to external and internal stimuli and facilitating a number of cellular structures such as lamellipodia or filopodia (Svitkina, 2018). These and other cellular structures are crucial for processes such as migration, cell division and phagocytosis. One of the main regulators of this dynamic actin network are the Rho GTPases. Rho GTPases are molecular switches that are ‘ON’ when they have bound GTP and ‘OFF’ when they have bound GDP (Bos et al., 2009). Guanine exchange factors (GEFs) facilitate the exchange of GDP to GTP, thereby turning the Rho GTPase on (Rossman et al., 2005). GTPase-activating proteins (GAP) activate the Rho GTPase’s intrinsic hydrolysation function, thereby turning the Rho GTPase off (Bos et al., 2009). The three most abundantly studied Rho GTPases are RhoA, Rac1 and Cdc42. RhoA signaling is associated with cell contraction and the formation of stress fibers (Hall, 1998). Rac1 induces lamellipodia, driven by actin polymerization and formation of a branched actin network (Hall, 1998). Cdc42 is inducing filopodia which contain parallel actin bundles (Hall, 1998).

The Rho GTPase activity is well defined in time but also in space (Pertz, 2010). For example, in a migrating cell RhoA activity is located at the rear edge and Cdc42 and Rac1 are active at the leading edge (Ridley, 2015). Another example of local Rho GTPase activity is the local and transient activity of Rho at a migration pore in an endothelial cell monolayer that is crossed by a leukocyte, to potentially confine the pore size and prevent leakiness of the vessel (Heemskerk et al., 2016; Mahlandt et al., 2021). Moreover, highly dynamic, subcellular Cdc42 and Rho alternating activity waves have been observed in a wound assay of the *Xenopus* oocyte, indicating cross talk between the Rho GTPases (Benink and Bement, 2005). These examples show that Rho GTPase activity is ideally monitored in living cells, with biosensors that can capture their activity over time but also the subcellular location of the activity. In the best case activity of multiple Rho GTPases would be monitored together in one cell to also measure cross talk (Welch et al., 2011).

A popular method to measure Rho GTPase activity in living cells is the use of FRET biosensors. These biosensors utilize Förster Resonance Energy Transfer (FRET), which can occur between two fluorescent proteins, typically cyan and yellow fluorescent proteins, when they come in close proximity. FRET biosensors are designed to undergo a conformational change altering the distance between the fluorescent proteins, upon presence of the target molecule, this can then be measured in a change of FRET ratio. Most Rho GTPase activity FRET sensors contain a Rho GTPase and a G-protein binding domain (GBD) which will bind the Rho GTPase in its GTP bound state, inducing the conformation change and thereby the FRET ratio change (Pertz, 2004). A number of FRET biosensors is available for Rho GTPase activity listed in the fluorescent biosensor database. By design, these FRET sensors are a measure of the GEF activity, because they contain the Rho GTPase. Some, but not all, lack the plasma membrane localization of the Rho GTPase. Plus, when the FRET signal is measured with a wide field microscope, these circumstances result in relatively low subcellular spatial resolution of the Rho GTPase activity detected with FRET sensors. Still, the ratiometric mode of imaging FRET sensors is beneficial for detection of gradients or activity in 3D imaging (Ponsioen et al., 2021).

Besides FRET sensors, another type of biosensor is available to detect Rho GTPase activity, namely localization-based biosensors. A localization-based Rho GTPase sensor binds to the endogenous Rho GTPase at its native location in the cell. In its inactive state this sensor will localize in the cytosol and relocalize to the endogenous Rho GTPase once its active, this results in a detectable fluorescence intensity change. This biosensor requires only a single fluorescent protein fused to the specific binding domain, thereby providing more spectral flexibility for multiplexing biosensors. However, this type of biosensor heavily relies on a specific Rho GTPase binding domain (GBD) for the active state, that has sufficient binding affinity.

We recently optimized a localization-based sensor for Rho (Mahlandt et al., 2021). This study is a continuation of that work, characterizing Cdc42 and Rac localization-based sensors. Cdc42 activity has been detected in *Xenopus* oocytes with the wGBD location sensor (Benink and Bement, 2005). Rac activity has been detected with location-based sensors using ABI1 (Innocenti et al., 2004), p67phox (Graessl et al., 2017), PBD (Petrie et al., 2012). However, their contrast is low in a cell culture model, requiring TIRF microscopy or pretreatment of the cells to round them up or treatment with nocodazole. Plus, for some of them their specificity for Cdc42 or Rac is poorly documented. These issues limit the usability of Cdc42 and Rac localization-based sensors.

Here we attempt to improve Cdc42 and Rac localization-based sensors for their relocation efficiency and specificity. Therefore, we select GBD candidates for Cdc42 and Rac specific localization-based biosensors from the recent mass spectrometry screen from Gillingham and colleagues that identify specific binders of active Rac and Cdc42 that will only bind on of the Rho GTPases in the active state (Gillingham et al., 2019). Additionally, known Rac and Cdc42 localization-based sensors were taken along. Candidates were subsequently evaluated for Rho GTPase binding in a microscopy-based analysis and the Rho GBDs were identified. The GBD candidates were tested for Rho GTPase specificity and location efficiency and the location efficiency was improved with a multi-domain approach.

The only candidate for a Rac1 location sensor that showed relocalization was CYRI-A, but the relocalization efficiency remains to be optimized. The Cdc42 localization-based sensor with the highest relocation efficiency was dimericTomato-WASp(CRIB), this GBD is also used in the original wGBD sensor (Benink and Bement, 2005). However, the most Cdc42 specific domain was N-WASP(CRIB). The Cdc42 localization-based sensor dimericTomato-WASp(CRIB) was multiplexed with the Rho localization-based sensor mTurquoise2-x3rGBD in a cell culture model, showing mutually exclusive activity.

## Results

### Candidate selection for location-based Cdc42 and Rac activity sensors

Designing a single-color biosensor indicating Rho GTPase activity by localization, requires a specific Rho GTPase binding domain (GBD) with high affinity for the GTP-bound Rho GTPase. The dynamic range and the specificity of the location biosensor will depend on the properties of the GBD. To find suitable GBDs, we re-analyzed the data from a mass spectrometry screen for Rho GTPase interacting proteins based on mitochondrial relocalization and proximity biotinylation (Gillingham et al., 2019). We looked first into GBDs for Rho, to compare the results of the mass spectrometry screen with the results of our cell-based assays (Mahlandt et al., 2021). High scoring proteins for interacting with constitutively active RhoA(Q63L) included ANLN part of the AniRBD Rho location sensor (Piekny and Glotzer, 2000), PKN1 part of aRho FRET sensor (Yoshizaki et al., 2003) and RTKN part of the rGBD Rho location sensor (Benink and Bement, 2005; Mahlandt et al., 2021) (**Fig. 1A,B**). This suggested that proteins with a high score in the mass spectrometry screen are potentially suitable as Rho GTPase activity biosensor. Indeed, the GBDs used for Cdc42 location sensors from, PAK1 used in the PBD location sensor (Itoh et al., 2002; Petrie et al., 2012) and N-WASP similar to WASp used in the wGBD location sensor (Benink and Bement, 2005) showed a high score in the screen (**Fig. 1A,B**). Interestingly, WASp itself did not show up in the screen. ABI1 which is used for Rac1 location sensors (Innocenti et al., 2004), also showed a high score in the screen (**Fig. 1A,B**). To create improved localization-based biosensors for Cdc42 and Rac, candidate GBDs were selected from top 30 scores of the mass spectrometry screen, that were specific for one Rho GTPase and their DNA was available on addgene (**Fig. 1A,B**). Additionally, known localization-based Rac and Cdc42 sensors were included. Plus, Bem1 and Bem3 a recommendation by the Laan lab for Cdc42 binding (Laan et al., 2015) and CYRI-A which was reported by Machesky and colleagues as a Rac binding protein (Le et al., 2021).

**Figure 1.**
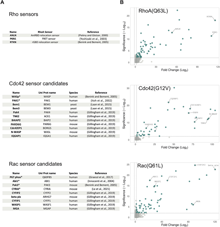
Candidate selection of Cdc42 and Rac GBDs for a localization-based biosensor. (**A**) Tables with candidate proteins. The asterisks mark candidates already used as biosensor. (**B**) Volcano plots adapted from Gillingham et al. 2019 for RhoA, Cdc42 and Rac1 with mutations locking them in their GTP-bound, active state, compared to a background of 17 GTPases locked in their inactive state. Significance on the Y axis is represented by log Student’s t-test p value and fold change on the X axis is represented by the root WD score. Blue color indicates hits above the 1.5-fold change threshold. Names of proteins selected for this study are indicated.

### Initial screen for Rho GTPase and sensor candidate colocalization

We have previously used nuclear localized, constitutive active Rho GTPases, but these are not accessible for larger proteins that cannot enter the nucleus. To initially assess the ability of the candidate proteins to localize with active Rho GTPases, they were tested for colocalization with mitochondrial tagged constitutively active Cdc42 or Rac1. To this end, a fusion with TOMM20 was used for mitochondrial localization. The candidate proteins were fluorescently tagged with mScarlet-I or mCherry and co-expressed with TOMM20-mTurquoise2-Cdc42(G12V)ΔCaaX or TOMM20-mTurquoise2-Rac1(G12V)ΔCaaX in HeLa cells. Most of the Cdc42 sensor candidates showed colocalization with the constitutively active Rho GTPases, where only three of the Rac sensor candidates showed colocalization (**Fig. 2A,B**). Pak1(CRIB) and WASp(CRIB), already used as localization-based sensors, only contained the GBD and showed a high level of colocalization. To test if the full length proteins were able to colocalize with the mitochondrial tagged constitutively active Rho GTPases, full length WASp was also tested for colocalization (**Fig. S1A**). It clearly colocalized and validated the approach. None of the candidates intrinsically localized at the mitochondria. Candidates colocalizing with the mitochondrial tagged Rho GTPase were further tested for their potential as localization-based sensors.

**Figure 2.**
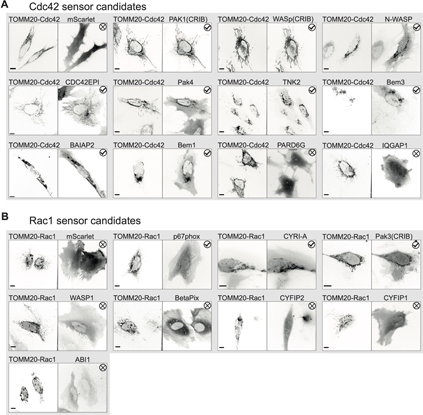
Initial screen for candidate colocalization with mitochondrial tagged constitutively active Rho GTPases. (**A**) Spinning disk microscopy images of HeLa cells co-expressing TOMM20-mTurquoise2-Cdc42(G12V)-ΔCaaX and the candidates tagged with mScarlet-I as indicated in image titles or mScarlet-I solely as a control. Check marks indicate colocalization and crosses no colocalization. Scale bars: 10 µm. (B) Spinning disk microscopy images of HeLa cells co-expressing TOMM20-mTurquoise2-Rac1(G12V)-ΔCaaX and either p67phox-mCherry, mCherry-CYRI-A, EGFP-Pak3(CRIB), mScarlet-I-WASP1, mCherry-BetaPix, mCherry-CYFIP2, mScarlet-I-CYFIP1, mScarlet-I-ABI1 or mScarlet-I solely as a control. Check marks indicate colocalization and crosses no colocalization. Scale bars: 10 µm.

### Identification and characterization of CRIB motifs in Cdc42 sensor candidates

Since the established Rho localization-based sensors utilize only the GBD and not the full length protein, we tried to identify the GBDs in the candidate proteins (Benink and Bement, 2005; Mahlandt et al., 2021; Piekny and Glotzer, 2000). For the Cdc42 biosensor candidates, the Cdc42/Rac interactive binding motif (CRIB), identified by the consensus sequence [I-S-x-P] (Burbelo et al., 1995; Manser et al., 1994), was found in a number of candidate proteins (**Fig. 3A**). No such consensus sequence could be found among the Rac sensor candidates. The Rac-p67phox crystal structure revealed CRIB independent binding, instead a tetratricopeptide repeat (TPR) domain facilitates the Rac binding (Lapouge et al., 2000). However, the p67phox Rac binding TPR domain was not found in the other Rac sensor candidates. The identified CRIB motifs for Cdc42 sensor candidates were fused with mScarlet-I. These m-Scarlet-I CRIB motifs for different Cdc42 sensor candidates were tested for colocalization with nuclear tagged constitutively activate Cdc42 (**Fig. 3B,C**). The influence of the expression level of the nuclear constitutively active Cdc42, and its nuclear enrichment, as well as the expression level of the sensor candidate on the nuclear enrichment of the sensor candidate were tested for correlation and found neglectable (**Figure S2A,B,C**). Interestingly, some candidate CRIB motifs were nuclear excluded, such as Bem3(CRIB2), Bem1(CRIB), and CDC42EPI(CRIB2) and we cannot exclude that these would be viable Cdc42 sensor candidates. On the other hand, Pard6g(CRIB) was nuclear enriched in control conditions. CDC42EPI(CRIB1+2), TNK2(CRIB), Pak4(CRIB), N-WASP(CRIB), WASp(CRIB) and Pak1(CRIB) were chosen for further testing because of their nuclear enrichment, when co-expressed with nuclear tagged constitutively active Cdc42 (**Fig. 3B,C**).

**Figure 3.**
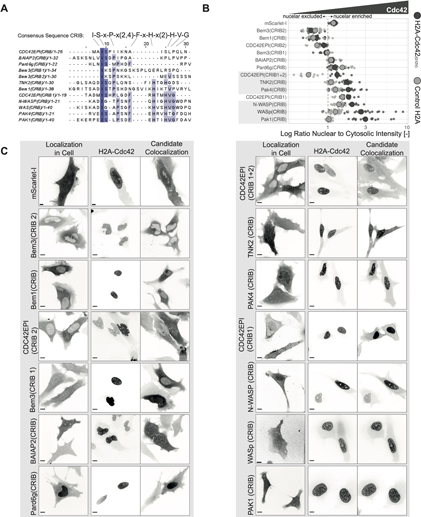
Identification and Cdc42 affinity characterization of Cdc42 sensor candidate CRIB motifs. (**A**) Amino acid sequence alignment for the indicated CRIB candidates with MUSCLE, CRIB consensus sequence shown above the alignment. Darker blue colors indicate higher conservation among the candidates. (**B**) Cdc42 affinity of the Cdc42 sensor candidate CRIBs measured by the ratio of nuclear to cytosolic intensity for the candidates fused to mScarlet-I, indicated in figure, in HeLa cells co-expressed with either H2A-mTurquoise2-Cdc42(G12V)-ΔCaaX or as a control H2A-mTurquoise2. Grey dashed line represents a ratio of 1. Small dots represent measurements from single cells. Larger grey dots represent the median. The number of cells analyzed is: BAIAP2(CRIB)-CDC42=16, BAIAP2(CRIB)-control=17, Bem1(CRIB)-Cdc42=20, Bem1(CRIB)-control=20, Bem3(CRIB1)-Cdc42=12, Bem3(CRIB1)-control=18, Bem3(CRIB2)-Cdc42=12, Bem3(CRIB2)-control=20, CDC42EPI(CRIB1)-Cdc42=10, CDC42EPI(CRIB1)-control=11, CDC42EPI(CRIB1+2)-Cdc42=9, CDC42EPI(CRIB1+2)-control=11, CDC42EPI(CRIB2)-Cdc42=10, CDC42EPI(CRIB2)-control=11, mScarlet-I-Cdc42=22, mScarlet-I-control=19, N-WASP(CRIB)-Cdc42=13, N-WASP(CRIB)-control=20, Pak1(CRIB)-Cdc42=15, Pak1(CRIB)-control=20, Pak4(CRIB)-Cdc42=16, Pak4(CRIB)-control=20, Pard6g(CRIB)-Cdc42=20, Pard6g(CRIB)-control=12, TNK2(CRIB)-Cdc42=18, TNK2(CRIB)-control=20, WASp(CRIB)-Cdc42=26, WASp(CRIB)-control=28 (**C**) Spinning disk microscopy images of HeLa cells expressing the Cdc42 sensor candidate CRIBs tagged with mScarlet-I (left panel) an HeLa cells co-expressing H2A-mTurquoise2-Cdc42(G12V)-ΔCaaX (middle panel) and mScarlet-I tagged Cdc42 sensor candidate CRIBs (right panel). Candidate names are indicated in the figure. Scale bars: 10 µm.

### Directly comparing Rac and Cdc42 affinity for CRIB Cdc42 sensor candidates

For the biosensor it is desirable to use a GBD that is specific for one Rho GTPase. The GBDs identified for the Cdc42 sensor candidates are all CRIB domains. As the motif name CRIB already suggests, these motifs may have binding affinity for Cdc42 as well as for Rac. However, they seem to have a preference for Cdc42 (Symons et al., 1996). Therefore, the six selected Cdc42 sensor candidate CRIBs were tested for their binding affinity to constitutively active Rac1 and Cdc42 in direct competition in a cellular environment. To this end, TOMM20-mTurquoise2-Cdc42(G12V)-ΔCaaX and H2A-mTurquoise2-Rac1(G12V)-ΔCaaX were co-expressed with the mScarlet-I tagged Cdc42 sensor candidate CRIBs. Hence, the sensor candidate can freely partition between Rac and Cdc42 binding. All six sensor candidates showed a clear preference for Cdc42 as they seemed to exclusively colocalize with the mitochondrial tagged constitutively active Cdc42, even though, judged by nuclear intensity, there was more constitutively active Rac1 expressed (**Fig. 4A**). However, in the absence of Cdc42 some of the Cdc42 sensor candidate CRIBs, namely of Pak1, WASp and Pak4, were able to bind constitutively active Rac1 tagged to the nucleus, while N-WASP(CRIB), TNK2(CRIB) and CDC42EPI(CRIB1+2) did not (**Fig. 4B**).

**Figure 4.**
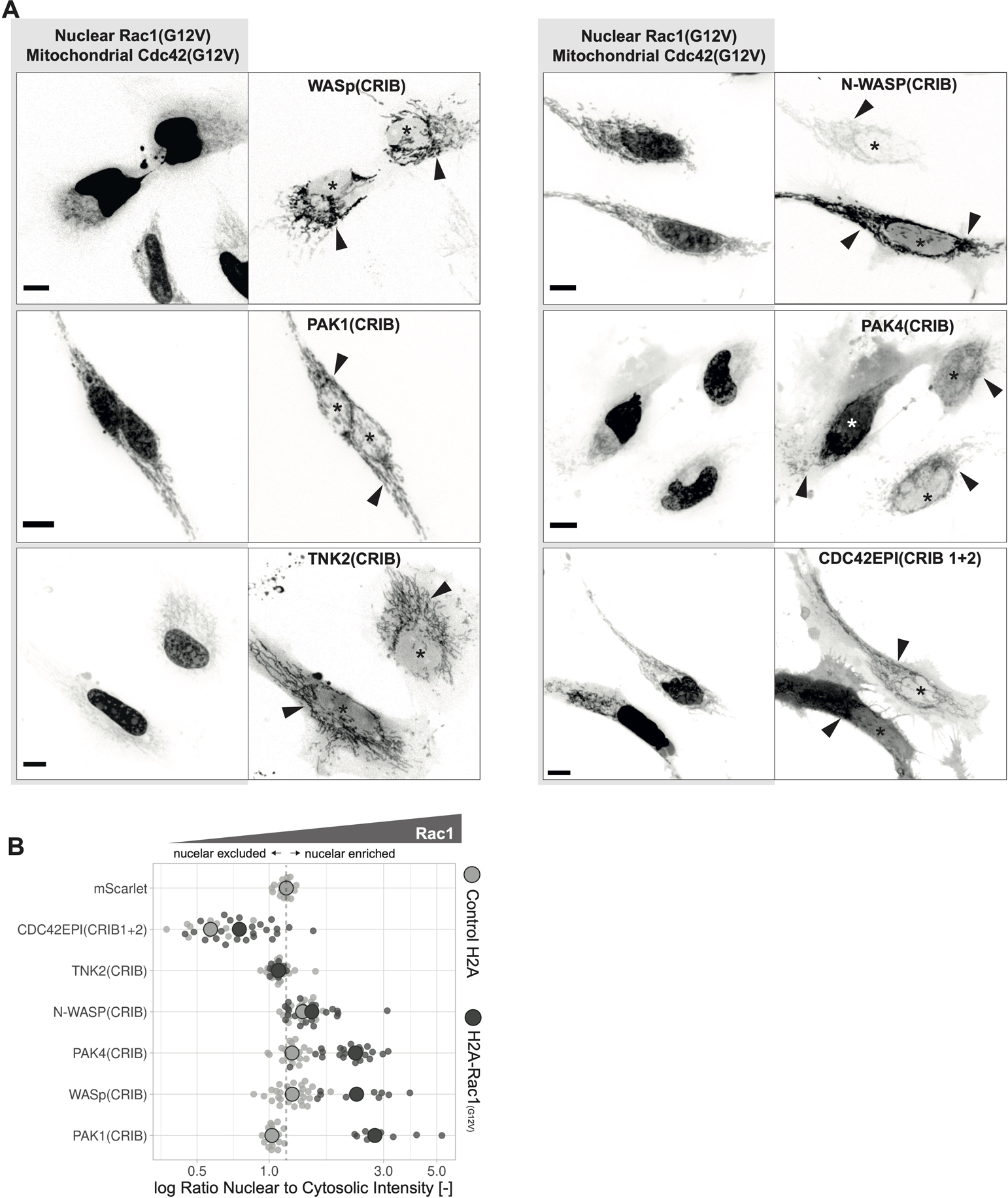
Comparison of Cdc42 sensor candidates CRIB’s affinity for Cdc42 and Rac1. (A) Maximum intensity projections of spinning disk images of HeLa cells co-expressing TOMM20-mTurquoise2-Cdc42(G12V)-ΔCaaX, H2A-mTurquoise2-Rac1(G12V)-ΔCaaX and mScarlet-I tagged WASp(CRIB), Pak1(CRIB), TNK2(CRIB), N-WASP(CRIB), Pak4(CRIB) and CDc42EPI(CRIB1+2). Asterisks indicate nucleus and arrows indicate mitochondria. Scale bars: 10 µm. (**B**) Rac1 affinity of the Cdc42 sensor candidate CRIBs measured by the ratio of nuclear to cytosolic intensity for the candidates fused to mScarlet-I, indicated in figure, in HeLa cells co-expressed with either H2A-mTurquoise2-Rac1(G12V)-ΔCaaX or as a control H2A-mTurquoise2. Grey dashed line represents a ratio of 1. Small dots represent measurements from single cells. Larger grey dots represent the median. The number of cells analyzed is: CDC42EPI(CRIB1+2)-control=11, CDC42EPI(CRIB1+2)-Rac1=25, mScarlet-I-control=19, N-WASP(CRIB)-control=20, N-WASP(CRIB)-Rac1=23, Pak1(CRIB)-control=20, Pak1(CRIB)-Rac1=10, Pak4(CRIB)-control=20, Pak4(CRIB)-Rac1=23, TNK2(CRIB)-control=20, TNK2(CRIB)-Rac1=12, WASp(CRIB)-control=28, WASp(CRIB)-Rac1=11. Control data is also shown in Fig. 3B.

### Optimizing Cdc42 sensor candidates for relocation efficiency

The ability of a sensor candidate to detect endogenous, active Cdc42 is critical for its use. Therefore, we evaluated the relocation efficiency of sensor candidate CRIBs in a synthetic system that can be used to activate endogenous Cdc42 in cells. The system used rapamycin induced heterodimerization of the two domains FRB and FKBP to recruit the DHPH domain of the Cdc42 specific GEF ITSN1 to the plasma membrane, where it induces activity of the endogenous Cdc42 (Inoue et al., 2005). The relocalization efficiency was tested in HeLa cells expressing Lck-mTurquoise2-FRB, YFP-FKBP-ITSN1(DHPH) and the Cdc42 sensor candidate CRIBs (**Fig. 5A, Movie 1**). The location sensor intensity should increase at the plasma membrane, which is the location of active Cdc42, and decrease in the cytosol. mScarlet-I-1xWASp(CRIB) showed that effect (**Fig. 5B,** **Movie 2,3**) and in comparison to the other candidates it showed the highest relocalization efficiency (**Fig. 5C**). To increase relocation efficiency, multiple CRIBs per sensor unit were tested, achieved either by sandwiching mScarlet-I between two CRIB domains or by the strong dimer fluorescent protein dimericTomato. One other strategy, namely cloning two WASp(CRIB) domains in tandem, directly after each other, lead to pre localization of the sensor at the plasma membrane (**Fig. 5B**) and was therefore not further tested for the other CRIBs. For the WASp(CRIB) the double CRIB domain as sandwich as well as the dimer with dimericTomato improved the relocalization efficiency as inferred from visual inspection (**Fig. 5A,B**).

**Figure 5.**
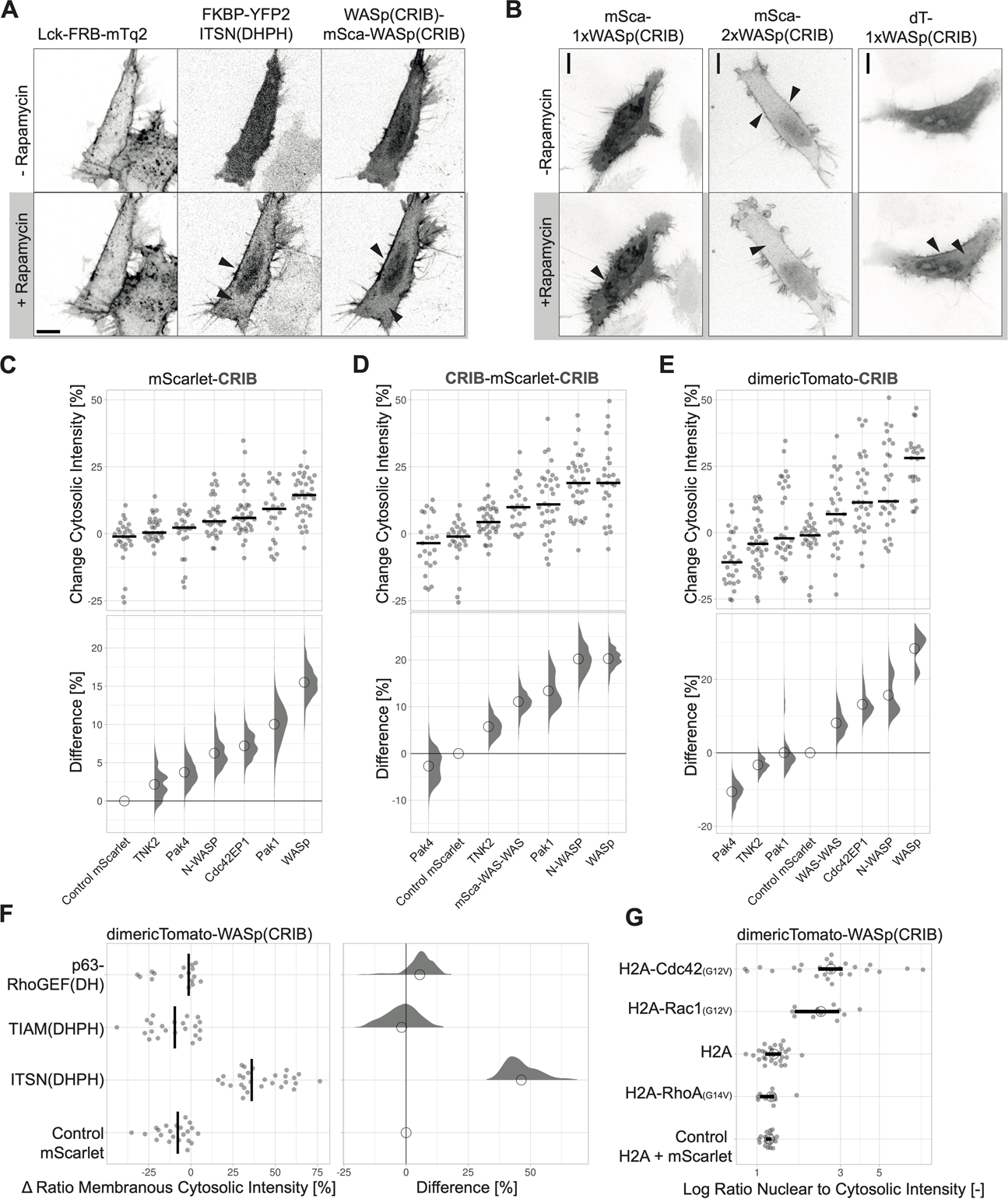
Optimizing relocation efficiency and characterizing Cdc42 sensor candidates. (**A**) Spinning disk microscopy images of a HeLa cell co-expressing Lck-FRB-mTurquoise2 (left), YFP-FKBP-ITSN1(DHPH) (middle) and WASp(CRIB)-mScarlet-I-WASp(CRIB) before and after the stimulation with 100 nM rapamycin. Arrows indicate intensity increase at the plasma membrane and depletion in the cytosol. Scale bar: 10 µm. (**B**) Spinning disk microscopy images of a HeLa cell co-expressing Lck-FRB-mTurquoise2 (not shown), YFP-FKBP-ITSN1(DHPH) (not shown) and either mScarlet-I-WASp(CRIB), mScarlet-I-WASp(CRIB)-WASp(CRIB) or dimericTomato-WASp(CRIB), before and after the stimulation with 100 nM rapamycin. Arrows indicate intensity increase at the plasma membrane and depletion in the cytosol. Scale bars: 10 µm. (**C,D,E**) Relocation efficiency of Cdc42 sensor candidates represented by the intensity change in the cytosol. (C) CRIBs fused to mScarlet-I, (D) mScarlet-I sandwiched between two CRIBs, and (E) CRIBs fused to dimericTomato. The relocation assay was performed as in (A). For the plot at the top, the median is represented as a black line and each dot represents the measurement of a single cell. The plot in the bottom shows the effect size relative to the mScarlet-I control. The bootstrap samples that are used to calculate the 95%CI of the effect size are shown as a distribution. The circle indicates the median. (F) Relocation of the dimericTomato-WASp(CRIB) sensor represented by the change in ratio of membranous over cytosolic intensity measured in HeLa cells expressing Lck-FRB-mTurquoise2, dimericTomato-WASp(CRIB) or mScarlet-I as a control and either YFP-FKBP-ITSN1(DHPH), YFP-FKBP-TIAM1(DHPH) or YFP-FKBP-RhoGEFp63(DH). FKBP-GEF recruitment was stimulated with 100 nM rapamycin. For the plot at the left, the median is represented as a black line and each dot represents the measurement of a single cell. The plot at the right shows the effect size relative to the mScarlet-I control. The bootstrap samples that are used to calculate the 95% confidence interval of the effect size are shown as a distribution. The circle indicates the median. (G) Rho GTPase specificity of for dimericTomato-WASp(CRIB) co-expressed in HeLa cells with either H2A-mTurquoise2-Cdc42(G12V)-ΔCaaX, H2A-mTurquoise2-Rac1(G12V)-ΔCaaX, H2A-mTurquoise2-RhoA(G14V)-ΔCaaX or H2A-mTurquoise2 as a control, represented by the ratio of nuclear over cytosolic sensor intensity. An additional control was the co-expression of mScarlet-I and H2A-mTurquoise2. The median is represented by a circle. Each dot represents the measurement of a single cell. The 95% confidence interval, calculated by bootstrapping, is indicated by a bar. The number of cells analyzed per conditions is: H2A-Cdc42=26, H2A mScarlet-I=19, H2A=28, H2A-Rac1=11, H2A-RhoA=11.

The Cdc42 sensor candidate CRIBs were compared for their relocalization efficiency as a single, domain, sandwich and dimer (**Fig. 5 C,D,E**). WASp(CRIB) showed the best recruitment efficiency as a single domain of roughly 15 %, which could be improved to roughly 20% with the sandwich and almost 30% with the dimericTomato. TNK2 and Pak4 CRIBs showed a low recruitment efficiency, in all three conditions. The N-WASP CRIB showed a relatively low recruitment efficiency as a single domain but this could be improved both, with the sandwich as well as the dimericTomato. The CDC42EPI(CRIB1+2)-mScarlet-I-CDC42EPI(CRIB1+2) sandwich was not generated successfully. Pak1 showed a relatively high recruitment efficiency as a single domain, which could be improved with the sandwich but not the dimericTomato construct. This maybe caused by the Ypet rest that was preserved during the protein engineering, which may not allow for the dimerization. Thus, N-WASP, CDC42EPI and Pak1 constructs had some relocation potential but dimericTomato-WASp(CRIB) showed the highest recruitment efficiency of almost 30%. Additionally, no correlation between absolute fluorescence intensity of the sensor and relocation efficiency could be measured (**Fig. S3A**).

The optimized dimericTomato-WASp(CRIB) Cdc42 relocation sensor was once more tested for its Cdc42 specificity. Therefore, TIAM and RhoGEFp63 were applied in the rapamycin assay to activate endogenous Rac and Rho, where before it was tested for the binding to overexpressed constitutively active Rho GTPases. The dimericTomato-WASp(CRIB) solely relocalized upon Cdc42 activation through ITSN recruitment (**Fig. 5F, Fig. S3B**). However, in the more artificial conditions of constitutively activate Rho GTPase tagged to the nucleus it is able to bind Rac1 but not RhoA (**Fig. 5G**).

### Comparing relocation efficiency for Rac sensor candidates

Based on the colocalization screen (figure 2B), we selected p67 phox, Pak3(CRIB), and CYRI-A as Rac sensor candidates. ABI1 was also included, as it was previously used as a Rac location sensor. Next, we assessed the Rac binding potential for the candidates, by screening for colocalization with the nuclear tagged constitutively active Rac1. Of the 4 candidates, p67 phox and Pak3(CRIB) did colocalize with Rac1 in the nucleus, and ABl1 and CYRI-A did not (**Fig. S4A**). Additionally, CYRI-A seemed to change the morphology of the nucleus and increase its size. Since the ABl1 fusion by itself is excluded from the nucleus, the assay with the constitutive Rac probe located in the mitochondria is more suitable.

Although CYRI-A localizes in the nucleus, it is not enriched when the active Rac1 is overexpressed. This may be explained by the putative myristoylation of CYRI-A (Le et al., 2021) which may be absent in the nuclear pool. Next, The rapamycin system was applied to activate endogenous Rac by the recruitment of TIAM to the plasma membrane, to measure relocation efficiency of the Rac sensor candidates (**Fig. 6 A,B, Fig. S4B**). TIRF microscopy was applied because the intensity changes were subtle and we and others noticed an improved signal to noise ratio for relocation sensors with TIRF microscopy (Graessl et al., 2017; Mahlandt et al., 2021). As the only Rac sensor candidate CYRI-A showed relocation to the plasma membrane upon Rac activation (**Fig. 6A,B**). In all conditions, we observed rapamycin induced recruitment of FKBP-TIAM to the plasma membrane and the TIRF field was relatively stable observed in the LCK-FRB-mTq2 channel, where the intensity only changed minimally (**Fig. S6B**). For Pak3(CRIB) we used a GFP-tagged version together with Lck-mTurquoise2-iLID and SspB-HaloTag-TIAM1(DHPH) for the activation of endogenous Rac. There was a non-Rac binding mutant available for CYRA-I, which indeed did not relocalize upon Rac activation (**Fig. S4C**). CYRI-A shows a relocation efficiency of roughly 30% imaged in TIRF mode and measured as the change in membranous intensity, but imaged in spinning disk mode and measured as the intensity depletion in the cytosol it is roughly 10% (**Fig. 6A, Fig. S6C**). To conclude, the best Rac sensor candidate is CYRA-I and its relocation efficiency could potentially still be optimized but that was not pursued in this study.

**Figure 6.**
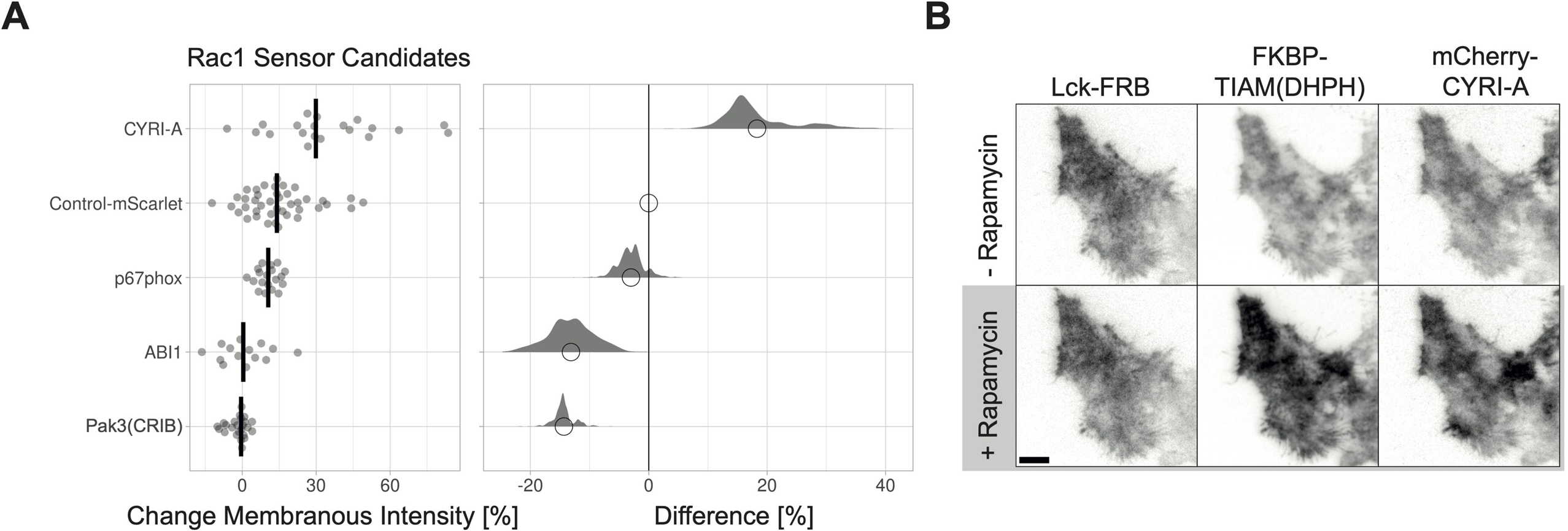
Comparison relocalization efficiency for Rac1 sensor candidates. (**A**) Relocation of the Rac sensor candidates represented by the change in membranous intensity measured in HeLa cells expressing Lck-FRB-mTurquoise2, YFP-FKBP-TIAM1(DHPH) and either mCherry-CYRI-A, mScarlet-I-ABI1 or p67phox-mCherry, stimulated with 100 nM rapamycin. Additionally, HeLa cells expressing Lck-mTurquoise-iLID, SspB-HaloTag-TIAM1(DHPH) and EGFP-Pak3(CRIB), stimulated with 440 nm laser light at 1% intensity for 1 s in 20 s intervals. For the plot at the left, the median is represented as a black line and each dot represents the measurement of a single cell. The plot at the right shows the effect size relative to the mScarlet-I control. The bootstrap samples that are used to calculate the 95%CI of the effect size are shown as a distribution. The circle indicates the median. The number of cells measured in two experiments based on independent transfections is: ABI1=14, Control-mScarlet=41, CYRI-A=20, p67phox=19, Pak3(CRIB)=24 (**B**) TIRF microscopy images for a HeLa cell expressing Lck-FRB-mTurquoise2 (left), YFP-FKBP-TIAM1(DHPH) (middle) and mCherry-CYRI-A (right) before and after stimulation with 100 nM rapamycin. Scale bar: 10 µm.

### Applying the optimized Cdc42 sensor in a multiplexing experiment and for localized activity

To apply the optimized Cdc42 relocation sensor dimericTomato-WASp(CRIB), a multiplexing experiment with the rGBD based Rho relocation sensor (Benink and Bement, 2005; Mahlandt et al., 2021) was performed. The Rho relocation sensor with the best relocation efficiency was also tagged with dimericTomato, namely dimericTomato2xrGBD. Therefore, different fluorescent tags were tested for the Rho sensor because it showed a better relocation efficiency than the dimericTomato-WASp(CRIB) sensor (Mahlandt et al., 2021). Other fluorescent proteins and the HaloTag were applied to the Rho relocation sensor (**Fig. S5A**). Of note, WASp(CRIB) can also be tagged with the HaloTag to allow for more flexibility in fluorescence color (**Fig. S5B**). For the Rho sensor, mTurquoise2 was chosen as a tag to avoid any crosstalk between the sensor signal especially with fraction of dimericTomato proteins that mature into a green chromophore (Strack et al., 2010). For the multiplex experiment, Hek cells expressing the Rho sensor mTurquoise2-3xrGBD and the Cdc42 sensor dimericTomato-WASp(CRIB), Lck-FRB-ECFP(W66A) non-fluorescent mutant and YFP—FKBP-ITSN1(DHPH) were first stimulated with human α-thrombin, activating endogenous Rho and subsequently with rapamycin activation endogenous Cdc42 via ITSN1 (**Fig. 7A,B, Fig. S5C,D,E, Movie 4**). Upon thrombin addition the Rho sensor relocalized to the plasma membrane and upon rapamycin addition the Cdc42 sensor relocalized to the plasma membrane, while the Rho sensor returned to the cytosol, indicating the inactivation of Rho at the moment of Cdc42 activation.

**Figure 7.**
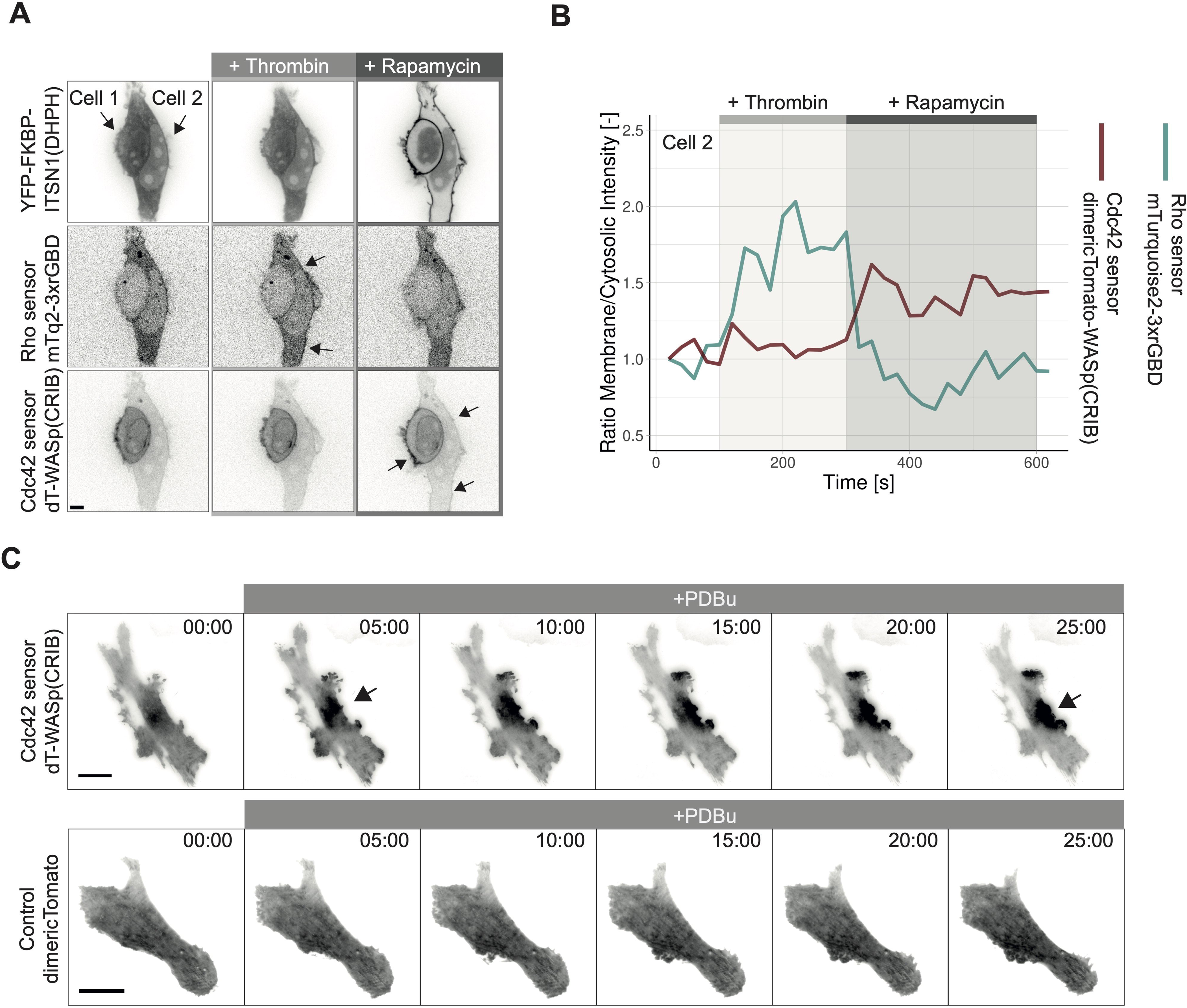
Applying the dimericTomato-WASp(CRIB) Cdc42 sensor for multiplexing and in a primary cells. (**A**) Spinning disk images of Hek 293T cells expressing Lck-FRB-ECFP(W66A)-black mutant, YFP-FKBP-ITSN1(DHPH) (top), Rho sensor mTurquoise2-3xrGBD (middle) and the Cdc42 sensor dimericTomato-WASp(CRIB) (bottom), first stimulated with 2 U/ml human α-thrombin, and then with 100 nM rapamycin. Each frame represents the maximum intensity projection of 3 frames over a time span of 1 min. Arrows in the top panel indicate the two cells. Arrows in the middle and bottom panel indicate intensity increase at the plasma membrane. Scale bar: 10 µm. (**B**) Time trace of the ratio membrane over cytosolic intensity for the in **B** depicted cell 2 for the Rho sensor mTurquoise2-3xrGBD (blue) and the Cdc42 sensor dimericTomato-WASp(CRIB). Thrombin addition is indicated by a light grey bar. Rapamycin addition is indicated by a dark grey bar. (**C**) TIRF microscopy images of a SUM159 cell expressing the Cdc42 sensor dimericTomato-WASp(CRIB) (upper panel) or dimericTomato as a control (lower panel), stimulated with PDBu (1 µM), as indicated by the grey bar. Arrows indicate the local signal accumulation of the Cdc42 sensor. Time is given in min:s from the beginning of the recording. Scale bar: 25 µm.

To test the ability of the sensor to reveal local, endogenous Cdc42 activity, the localization of the sensor at invadopodia was imaged. Invadopodia are F-actin rich, protrusive structures that enable cancer cells to invade tissue. Cdc42 has been shown to be involved in their assembly (Kreider-Letterman et al., 2023; Murphy and Courtneidge, 2011). The Cdc42 sensor dimericTomato-WASp(CRIB) accumulated in a spatially restricted manner upon invadopodia formation, induced by phorbol ester Phorbol 12,13-dibutyrate (PDBu) treatment of SUM159 breast cancer cells (**Fig. 7C, Fig. S5F, Movie 5, Movie 6**). No accumulation of a control, untagged dimericTomato, was observed under the same conditions (**Movie 7**). These results showcase the potential of the location-based sensor to visualize the subcellular location of endogenous Cdc42 activity.

## Discussion

Relocation based biosensors are powerful tools to visualize spatiotemporal aspects of Rho GTPase activity. However, relocation based sensors for Rac and Cdc42 are sparsely characterized and show poor performance in mammalian cell systems. Therefore, we examined GBDs from existing location-based sensors and new sensor candidate GBDs for their relocation biosensor potential. The best performing candidate biosensors were optimized and their selectivity and affinity were tested in cell-based assays. As a result, we identify dimericTomato-WASp(CRIB) as the most optimal sensor for detecting endogenous Cdc42 activity. A good performing Rac1 sensor has not yet been found, although we show that a recently identified Rac binding protein can be used to detect endogenous active Rac. Finally, we have made color variants of a Rho biosensor that we previously optimized to enable multiplex imaging of Rho and Cdc42.

Searching for a Cdc42 specific relocation sensor, dimericTomato-WASp(CRIB) showed the highest relocation efficiency and was Cdc42 specific in the GEF induced Rho GTPase activity assay but not in the constitutively active nuclear tagged Rho GTPase assay, meaning it is potentially able to bind constitutively active Rac1. However, the GEF induced Rho GTPases activity assay resembles the physiological conditions better than the constitutively active Rho GTPase assay. Therefore, our results suggest that the WASp(CRIB) will bind Cdc42 specifically under physiological conditions. Another Cdc42 sensor candidate, namely N-WASP(CRIB) was specific for Cdc42 in both assays and showed a high relocation efficiency. However, within the group of closely Cdc42 related Rho GTPases, namely Tc10, RHoT and Chp, N-WASP and WASp have no specificity and have been shown to bind all members (Takenawa and Suetsugu, 2007). The specificity for active GTP-bound Cdc42, in comparison to the GDP-bound form, has been shown for the WASp(CRIB), demonstrating that a CRIB domain can differentiate reliably between active and inactive Rho GTPases (Symons et al., 1996). Consequently, N-WASP(CRIB)-mScarlet-I-N-WASP(CRIB) is the recommended choice of sensor when Cdc42 specificity is crucial but if a higher relocation efficiency is necessary dimericTomato-WASp(CRIB) would be the sensor of choice to detect endogenous Cdc42 activity.

The search for Rac specific relocation sensors was less successful. We only found one candidate, namely CYRI-A to localize to the plasma membrane upon GEF induced endogenous Rac activation. However, the relocation efficiency was low and remains to be optimized. The other candidates that had previously been used as Rac sensor did not show localization in our experiments. For example, ABI1 has previously been used in cell that were rounded up before imaging and thereby creating an optimal cell shape to detect relocalization (de Beco et al., 2018). In another study p67phox relocation has been imaged in cells treated with nocodazole and imaged with TIRF microscopy, which improves the signal to noise ratio for relocation sensors (Graessl et al., 2017). That study used U2OS cells which might have higher endogenous Rac levels, which increases the relocalization efficiency. In our experimental conditions the relocation efficiency of this sensor was probably too low to be detected. Besides the endogenous Rac levels, the expression level of the sensor also plays a role, since the superfluous fraction of the sensor occludes the relocalizing fraction. None of the Rac binding candidates from the mass spectrometry screen performed by Gillingham and colleagues colocalized with Rac in our assay. Their Rac binding might depend on the WAVE regulatory complex, as ABI1 and CYFIP1/2 are members of the WAVE regulatory complex (Gillingham et al., 2019; Takenawa and Suetsugu, 2007).

Selecting relocation sensor candidates from the proximity biotinylation mass spectrometry screen for G-protein interactors by Gillingham and colleagues, was only partly successful (Gillingham et al., 2019). A number of the selected sensor candidates did not colocalize with the constitutively active Rho GTPase, especially for Rac1. That was surprising because that mass spectrometry screen also applies mitochondrial tagged constitutively active Rho GTPases. However, the readouts are different, where we screened for colocalization the mass spectrometry screen measured proximity biotinylation, which might be more sensitive. Additionally, we overexpressed both the Rho GTPase and the candidate, whereas the mass spectrometry screen only overexpressed the Rho GTPase. It is possible that the superfluous overexpressed candidate protein concealed the binding fraction. If the candidates bind the Rho GTPase in a complex, the endogenous binding partners should have been present in both cases, but the mass spectrometry screen used Hek293T cells and here HeLa cells were used, which might have different expression levels for binding partners.

The ideal method to identify a GBD for a relocation sensor remains unclear. Apparently, a high score in the proximity biotinylation GTPase interactor mass spectrometry screen was not sufficient for an efficient relocation sensor (Gillingham et al., 2019). The ideal GBD for a sensor requires a strong enough binding affinity and the specificity for one active Rho GTPase. The selection of candidates could potentially be improved be screening for binding affinity to the active Rho GTPase by determining a K_D_. A high WD score in the mass spectrometry screen does not predict the relocation efficiency of the sensor candidates. For example, N-WASP and TNK2 score very similarly in the mass spectrometry screen but TNK2 shows almost no relocation, whereas N-WASP shows high relocation efficiency. Also, previously tested PKN1, which scores high in the mass spectrometry screen for active RhoA, could not successfully be applied as a localization-based sensor (Mahlandt et al., 2021).

Also the function of the candidate proteins did not explain their relocation sensor potential. CYFIP1, CYFIP2 and ABI1 are part of the WAVE regulatory complex and CYRI-A is involved in its regulation (Rottner et al., 2021). The direct Rac1 binding in the WAVE regulatory complex is mediated by SRA1 (CYFIP1/2 orthologs) and IRSp53. ABI1 does not seem to have a direct interaction with Rac1. However, neither ABI1 nor CYFIP1/2 show colocalization with constitutively active Rac1 localized at the mitochondria. This might suggest that the entire WAVE complex is required for Rac1 binding including the synergistic binding to the plasma membrane. CYRI-A is not part of the WAVE complex but it can disrupt the Rac1-WAVE regulatory complex. CYRI-A has been shown to indicate Rac1 activity by its localization (Le et al., 2021), which confirms the findings of our study. Pak1, Pak3, Pak4, PAR6A and TNK2 are kinases (Bishop and Hall, 2000; Garrard et al., 2003), N-WASP, WASp, IQGAP1, p67phox, BAIAP2, CDD42EPI and Bem1 are scaffold proteins (Bishop and Hall, 2000; Burbelo et al., 1999; Laan et al., 2015). Bem3 is a GAP (Zheng et al., 1993), MGA is a transcription factor (Hurlin et al., 1999) and BetaPix is a Rac1 GEF (Welch et al., 2011). However, both scaffold proteins and kinases contained candidates that showed relocation and others did not even colocalize with the constitutively active Rho GTPase. Consequently, protein function does not determine the potential of a GBD for a localization-based sensor. Neither the WD score in the mass spectrometry screen, nor the protein function allow a prediction of the GBD’s potential to relocalizes to an active Rho GTPase. Thus, it remains unclear which properties define a good GBD for a location-based sensor.

Another challenge is the Rho GTPase specificity of the relocation-based sensor. For example, Pak1(CRIB) was first used in a Rac1 FRET sensor (Kraynov et al., 2000). Next, Pak1(CRIB) has been utilized in Cdc42 FRET sensors and in an intensiometric Cdc42 sensor (Hanna et al., 2014; Itoh et al., 2002; Kim et al., 2019). However, Pak1(CRIB), also named PBD sensor, has then been reintroduced by Weiner and colleagues as a Rac1 specific location-based sensor and is often used in neutrophil HL60 cells (Brunetti et al., 2022; Graziano et al., 2019; Le et al., 2021; Weiner et al., 2007). We showed that Pak1(CRIB) can bind constitutively active Rac1 and Cdc42 but in direct competition it binds Cdc42. That strongly contrasts a study claiming Pak1(CRIB) colocalizes with active Rac1 but not Cdc42 (Srinivasan et al., 2003). That study used the plasma membrane localization of the overexpressed active Rho GTPase, which is not as distinct as the nucleus or the mitochondria, therefore the colocalization might be challenging to assess. The nuclear or mitochondrial tagged constitutively active Rho GTPases that we have generated provide a clearer read out and allow direct comparison of Rho GTPase binding in a cell-based assay. Also, Itoh and colleagues find the Pak PBD to perform better in the Cdc42 FRET sensor than in the Rac1 sensor (Itoh et al., 2002). We conclude that Pak1(CRIB) is binding both Rac1 and Cdc42, with a preference for Cdc42, and cannot be applied as a specific relocation sensor for either of them. However, it was possible to achieve Rho GTPase specificity with a GBD domain as we showed for the CRIBs of N-WASP, TNK2 and CDC42EPI. Yang and colleagues confirm the Cdc42 specificity of TNK2 (Yang and Cerione, 1997).

The common amino acid sequences recognizing Cdc42 and Rac in these proteins is the CRIB motif. This motif contains the consensus sequence I-S-x-P-x(2,4)-F-x-H-x(2)-H-V-G (Burbelo et al., 1995). The name suggest binding of Rac and Cdc42 with preference for Cdc42 (Symons et al., 1996). A somewhat larger amino acid sequence, of about 30 amino acids, is necessary to bind Cdc42 or Rac1 as was shown at the example of Pak1(CRIB) (Thompson et al., 1998). We also used a region of about 30 amino acids including the CRIB and could confirm the preference of binding Cdc42. In contrast, P67phox achieves specific Rac binding with a TPR domain and two amino acids, namely Ala-27 and Gly-30 found in Rac1 but not in Cdc42, seem to be sufficient for specific Rac1 binding of p67phox (Lapouge et al., 2000). Thus, Rac1 or Cdc42 specificity is possible and can be facilitated by a few amino acids. However, concentrations also play a role in the binding affinity we see all CRIB candidates exclusively binding active Cdc42 in direct competition with Rac1, but if only active Rac1 is available they bind Rac1.

To increase the relocation efficiency, the multiple binding domain strategy worked well for the rGBD Rho sensor, for PH domain based lipid sensor and a Ras activity sensor (Augsten et al., 2006; Goulden et al., 2019; Mahlandt et al., 2021). However, it was only partly successful in increasing the relocation efficient of the CRIB domain based sensors. Two WASp(CRIB) domains in tandem resulted in prelocalization of the construct at the plasma membrane. This could be explained if the domain contained parts with plasma membrane affinity, which could increase the binding to the plasma membrane in a multi-domain construct. For example, WASP, N-WASP have been shown to interact with phosphoinositides (Takenawa and Suetsugu, 2007). Arranging the domains with more space in between by creating a WASp(CRIB)-mScarlet-I-WASp(CRIB) sandwich or dimers of dimericTomato-WASp(CRIB) partly prevented the prelocalization and increased the relocation efficiency. However, this approach did not increase the relocation efficiency in the same manner for all tested CRIBs.

To reliably detected relocalization the following points should be taken in account. A low expression level is optimal to observe relocalization of the sensor, because the larger the fraction of sensor molecules that bind the limited endogenous Rho GTPase, the higher the relocalization contrast. However, the lower the fluorescent intensity the worse the signal to noise ratio in the images. Hence, the expression level needs to be optimized between signal to noise ratio and relocation contrast. To find the optimal expression level the low expressing CMVdel promoter can be applied (Watanabe and Mitchison, 2002) and cell sorting could be used. Also, advanced microscopy techniques can improve the signal to noise ratio, enabling the imaging of a lower fluorescent signal. We noticed that spinning disk microscopy and TIRF microscopy were beneficial for the imaging of relocation sensor, as did others (Graessl et al., 2017). Additionally, it is essential to be aware of the fact that morphology changes such as local protrusions, especially in ventral orientation, can cause intensity increases that are not related to relocation (Dewitt et al., 2009; van der Linden et al., 2021). That issue could be addressed with a reference signal for the plasma membrane intensity (Mahlandt et al., 2021).

Location-based Rho GTPase activity sensor give the opportunity to image the precise location and timing of endogenous Rho GTPase activity. In comparison, the common ratiometric FRET sensor do provide a semi-quantitative ratio metric readout, but they are not detecting the endogenous Rho GTPase but rather the activity of endogenous GEFs and GAPs. We showed high spatial resolution in endogenous Cdc42 activity in invadopodia in this study and another (Kreider-Letterman et al., 2023). Furthermore, single-color location based sensors are a valuable tool in optogenetics because the photo-activation spectrum overlaps with the spectrum of the common CFP-YFP FRET pair. Moreover, multiplex imaging of biosensors is a powerful approach to unravel Rho GTPase crosstalk and single-color, location based sensor are of advantage for this because they simply occupy a smaller part of the light spectrum. We have shown the multiplexing of the optimized Rho and Cdc42 activity location-based sensor in a cell culture model, indicating mutually exclusive activity. Previously this multiplexing of location-based sensors has been achieved in the *Xenopus* oocyte model (Benink and Bement, 2005; Moe et al., 2021) and in nocodazole treated U2OS cells (Graessl et al., 2017).

In conclusion, we characterized and optimized single-color, genetically encoded, location-based biosensors for the endogenous Cdc42 or Rac activity. This will improve detection of active Cdc42 or Rac, especially in optogentics and multiplex experiments and will thereby contribute to a better understanding of Rho GTPase signaling. Additionally, the tools that we developed for characterization, namely localized constitutively active Rho GTPases, will be beneficial for future studies that identify and validate Rho GTPase relocation sensors.

## Methods

### Plasmids and Cloning

The BEM 1 and BEM3 containing plasmids were a gift from Lidewij Laan. The PAK4 containing plasmid was a gift from William Hahn & David Root (addgene # 23713). The TNK2 containing plasmid was a gift from William Hahn & Jean Zhao (addgene #20654), this construct contains the first 499 amino acids of TNK2. The BAIAP2 wt containing plasmid was a gift from Anne Brunet (addgene #31656). The Par6A containing plasmid was a gift from Ian Macara (addgene #15472). The CDC42EP1 WT containing plasmid was a gift from Anne Brunet (addgene #69819). The GFP-wGBD, in this study WASp(CRIB), containing plasmid was a gift from William Bement (addgene #26734). The Ypet-PBD, in this study Pak1(CRIB), containing plasmid was a gift from Kenneth Yamada (addgene #105290). The mEmerald-N-WASP containing plasmid was a gift from Michael Davidson (addgene #54199). The pEGFP-IQGAP1 containing plasmid was a gift from David Sacks (addgene #30112). The mCherry-p67phox containing plasmid was a gift from Perihan Nalbant. The pEGFP-ABI1 containing plasmid was a gift from Giorgio Scita (addgene #74905). The CYFIP2-mCherry containing plasmid was a gift from Josef Kittler (Addgene plasmid # 122052). The ECFP-betaPIXa containing plasmid was a gift from Rick Horwitz (addgene #15235). The CYFIP1-GFP containing plasmid was a gift from Josef Kittler (addgene #109139). The mEmerald-WASP1 containing plasmid was a gift from Michael Davidson (addgene #54314). The MGA containing plasmid was a gift from Guntram Suske (addgene #107715). The GFP-Pak3(CRIB) containing plasmid was a gift from William Bement (addgene #26735). The mCherry-CYRI-A and mCherry-CYRIA(49A) containing plasmids were a gift from Laura Machesky. The full length WASp containing plasmid was a gift from David Rawlings (addgene #36248).

### Tagging candidates with mScarlet-I

mScarlet-I, described previously (Bindels et al., 2016), was used to fluorescently tag the candidate proteins. All backbones were dephosphorylated after digestion. mScarlet-I-IQGAP, mScarlet-I-ABI1 and mScarlet-I-WASP1 were created by digesting the backbones pEGFP-IQGAP, pEGFP-ABI1 and mEmerald-WASP1 with AgeI and BsrGI and ligating them to the likewise digested insert mScarlet-I. mScarlet-I-Pak4 was created by PCR amplifying Pak4 (primers: FW-5’TATAGAATTCGATGTTTGGGAAGAGGAA3’, RV-5’TATAGGATCCTTA TCATCTGGTGCGGTTCT3’), digesting it with EcoRI and BamHI and ligating it to the likewise digested backbone mScralet-I-C1. mScarlet-I-BAIAP2 was created by digesting BAIAP2 with EcoRI and XbaI and ligating it to the likewise digested backbone mScarlet-I-C1. mScarlet-I-CDC42EPI was created by PCR amplifying CDC42EPI (primers: FW-5’TATAGAATTCGATGCCCGGCCCCCAG3’. RV-5’TATAGGATCCTTACACCTTGACCTCATCATC3’), digesting it with EcoRI and BamHI and ligating it to the likewise digested backbone mScralet-I-C1. mScarlet-I-MGA was created by PCR amplifying mScarlet-I (primers: FW-5’ATATAGCGCTATGGTGAGCAAGGGCG 3’, RV-5’ATATAGCGCTCCTTGTACAGCTCGTCCATG3’), digesting it with AfeI and ligating it to the likewise digested backbone MGA. mScarlet-I-MGA could not be expressed in HeLa cells. mScarlet-I-Pard6g was created by digesting Pard6g with BamHI and ligating it to the likewise digested backbone mScarlet-I-C1. mScarlet-I-Bem1 was created by PCR amplifying Bem1 (primers: FW-5’GCTGTACAAGTCCATGCTGAAAAACTTCAAACTCTCAA3’, RV-5’ATATCTCGAGTTAGAGTCTAATATCGTGAACGGAAATTT3’), digesting it with BsrGI and XhoI and ligating it to the likewise digested backbone mScarlet-I-C1. mScarlet-I-Bem3 was created by PCR amplifying Bem3 (primers: FW-5’ATATTCCGGAATGACAGATAATTTGACCACAACTC, RV-5’ATATGAATTCTCAAACCTGAGGAATATGTATATCTACTTT3’), digesting it with BspEI and EcoRI and ligating it to the likewise digested backbone mScarlet-I-C1. mScarlet-I-TNK2 was created by PCR amplifying TNK2 (primers: FW-5’ATATTGTACAAGATGCAGCCAGAGGAGGGCA, RV-5’ATATAAGCTTTCAGCGCTTGTGGTGG3’), digesting it with BsrGI and HindIII and ligating it to the likewise digested backbone mScarlet-I-C1. mScarlet-I-WASp was created by PCR amplifying WAS (primers: FW-5’ATATAAGCTTCGATGAGTGGGGGCCCAA, RV-5’ATATGGATCCTCAGTCATCCCATTCATCATCTTC), digested with HindIII and BamHI and ligated to the likewise digested backbone mScarlet-I-C1.

### Single mScarlet-I-CRIB constructs

CRIBs were identified with the ScanProsite tool (Expasy, SIB Swiss Institute of Bioinformatics) by searching for I-[SG]-x-P, a part of the full consensus sequence I-S-x-P-x(2,4)-F-x-H-x(2)-H-V-G (Burbelo et al., 1995). A region of roughly 300 base pairs was chosen to clone the identified CRIBs in a mScarlet-I-C1 backbone. mScarlet-I-BAIAP2(CRIB), mScarlet-I-Pard6g(CRIB), mScarlet-I-Bem1(CRIB), mScarlet-I-Pak4(CRIB), mScarlet-I-CDC42EPI(CRIB1), mScarlet-I-CDC42EPI(CRIB2), mScarlet-I-CDC42EPI(CRIB1+2), mScarlet-I-Bem3(CRIB1), mScarlet-I-Bem3(CRIB2) and mScarlet-I-N-WASP(CRIB) were created by PCR amplifying the CRIBs, digesting them with BsrGI and XhoI and ligating them to the likewise digested mScarlet-I-C1 backbone. BAIAP2(CRIB) was amplified with the primers: FW-5’ GCTGTACAAGTCCGAGCGCGCGGTGC3, and RV-5’ATACCTCGAGTTAGCCTGTGACGCCGTTC3’. Pard6g(CRIB) was amplified with the primers: FW-5’ GCTGTACAAGTCCAGCTTCGGAGCAGGCA3’ and RV-5’ACCTCGAGTTACACCCGCACGCG3’. Bem1(CRIB) was amplified with the primers: FW-5’GCTGTACAAGTCCATGCTGAAAAACTTCAAACTCTCAA3’ and RV-5’ATACCTCGAGTTATTCACCTTCCATGAAAGATAGTTCC3’. Pak4(CRIB) was amplified with the primers: FW-5’GCTGTACAAGTCCATGTTTGGGAAGAGGAAGAAGC3’ and RV-5’ATACCTCGAGTTACAGCAGCGTGAGGGC3’. CDC42EPI(CRIB1) was amplified with the primers: FW-5’ATATTGTACAAGATGATGCCCGGCCCCCAG3’ and RV-5’ACCTCGAGTTAGATGGCCGGTGGAGC3’. CDC42EPI(CRIB2) was amplified with the primers: FW-5’ATATTGTACAAGATGGCCATCTCCCCCATCATC3’and RV-5’ACCTCGAGTTAGAGAGAGTCAGAGCGGC3’. CDC42EPI(CRIB1+2) was amplified with the primers: FW-5’ ATATTGTACAAGATGATGCCCGGCCCCCAG3’ and RV-5’ACCTCGAGTTAGAGAGAGTCAGAGCGGC3’. Bem3(CRIB1) was amplified with the primers: FW-5’ATATTGTACAAGATGGCCTCTGTTACCTATACGACG 3’ and RV-5’ACCTCGAGTTAGGATGTTATAGAAGAATAAACGCGG3’. Bem3(CRIB2) was amplified with the primers: FW-5’ATATTGTACAAGATGATGAACCATATAGGTATTACAATTTCAAATGA3’ and RV-5’ACCTCGAGTTACCCGCTTAGTCTAAATATGCCT3’. N-WASP(CRIB) was amplified with the primers: FW-5’ ATATTGTACAAGATGATCTCCCACACCAAAGAAAAGAA3’ and RV-5’ACACCTCGAGTTATGGTGGTGCTTGTCTTCG3’. mScarlet-I-Pak1(CRIB) was created by digesting mScarlet-I with AgeI and BsrGI and ligating it to the likewise digested backbone YPet-PBD. A rest of the YPet sequence remained in the construct. mScarlet-I-TNK2(CRIB) was created by PCR amplifying TNK2(CRIB) with the primers FW-5’GCTGTACAAGTCCGTGGGGCCCTTCCCT3, and RV-5’ACAAGCTTTTAGTTTCCCAGATACAGTTCGTCAATCCTGTCCGG3’, digesting it with BsrGI and HindIII and ligating it to the likewise digested mScarlet-I-C1 backbone. The reduced expression GFP β-actin plasmid was a gift from Rick Horwitz and Tim Mitchison (addgene #31502). In this plasmid the base pairs 91–544 of the enhancer region in the CMV promoter are deleted; in the following called CMVdel. CMVdel-mScarlet-I-WASp(CRIB) was created by first digesting the backbone mCherry-wGBD (in this study named WASp(CRIB)) with AgeI and BsrGI and ligating the likewise digested mScarlet-I to it. Then mScarlet-I-WASp(CRIB) was excised with AgeI and MluI and ligated to the likewise digested CMVdel reduced expression GFP β-actin plasmid. CMVdel-mScarlet-I-2xWASp(CRIB) was created by PCR amplifying WASp(CRIB) (primers: FW-5’TATATATGTACATCCTCCCTAGCCCAGCTGATAAG3’ and RV-5’TATATAGGTACCCCCTCTCCTGGCGCCTCATCTC3), digesting it with BsrGI and Acc65I and ligating it to the backbone mScarlet-I-WASp(CRIB), which was digested with BsrGI.

### Sandwich CRIB-mScarlet-I-CRIB constructs

WASp(CRIB)-mScarlet-I-WASp(CRIB) was created by PCR amplifying WASp(CRIB) (primers: FW-5’ATATGCTAGCATGCCTAGCCCAGCTGATAAGAAAC and RV-5’ATATACCGGTGAGAACTCCTGGCGCCTC3’), digesting it with NheI and AgeI and ligating it to the likewise digested backbone mScarlet-I-WASp(CRIB). Pak1(CRIB)-mScarlet-I-Pak1(CRIB) was created by PCR amplifying Pak1(CRIB) (primers: FW5’ ATATGCTAGCATGGCTTCGAATTCCAATAAAAAGAAAGAG3’ and RV-5’ATATACCGGTGCTCTAGATCCGGTGGATCCAG3’), digesting it with NheI and AgeI and ligating it to the likewise digested backbone mScarlet-I-Pak1(CRIB). N-WASP(CRIB)-mScarlet-I-N-WASP(CRIB) was created by PCR amplifying N-WASP(CRIB) (primer: FW-5’ ATATGCTAGCATGATCTCCCACACCAAAGAAAA3, and RV-5’ ATATACCGGTGCTGGTGGTGCTTGTCTTCG), digesting it with NheI and AgeI and ligating it to the likewise digested backbone mScarlet-I-N-WASP(CRIB). Pak4(CRIB)-mScarlet-I-Pak4(CRIB) was created by PCR amplifying Pak4(CRIB) (primers: FW-5’ ATATGCTAGCATGTTTGGGAAGAGGAAGAAGC3’ and RV-5’ ATATACCGGTGCGAGGGGCTTGGGCC3’), digesting it with NheI and AgeI and ligating it to the likewise digested backbone mScarlet-I-Pak4(CRIB). TNK2(CRIB)-mScarlet-I-TNK2(CRIB) was created by PCR amplifying TNK2(CRIB) (primers: FW-5’ATATGCTAGCATGTCCGTGGGGCCCTTC and RV-5’ ATATACCGGTGCGTTTCCCAGATACAGTTCGTCA3’), digesting it with NheI and AgeI and ligating it to the likewise digested backbone mScarlet-I-TNK2(CRIB).

### Dimer dimericTomato-CRIB constructs

DimericTomato-WASp(CRIB), −2xWASp(CRIB), -Pak1(CRIB), -CDC42EPI(CRIB1+2), -N-WASP(CRIB), - Pak4(CRIB), -TNK2(CRIB) by digesting the corresponding mScarlet-I-CRIB backbones (described above) with AgeI and BsrGI and ligating the likewise digested insert dimericTomato to it.

### Location sensor color variants

DimericTomato-2xrGBD (addgene #129625), mNeonGreen-2xrGBD (addgene #129624) 3xmNeonGreen-3xrGBD (addgene #176101) were described previously (Mahlandt et al., 2021). The cysteine-free, or secretory HaloTag version 7 was provided by Ariana Tkachuk in consultation with Erik Snapp (Janelia Research Campus of the Howard Hughes Medical Institute). The HaloTag was PCR amplified (primers: FW 5’-ATATACCGGTCGCCACCATGGCCGAGATCGGCA-3’ and RV 5’-ATATTGTACACGCCGCTGATCTCCAGG-3’) and AgeI, BsrGI digested and ligated into a likewise digested C1 (Clontech) backbone, creating C1-HaloTag. mTurquoise2-3xrGBD, HaloTag-3xrGBD and 3xmTurquoise3-3xrGBD were created by digesting the backbone 3xmNeonGreen-3xrGBD with AgeI and BsrGI and ligating the likewise digested inserts, mTurquoise2, 3xmTurquoise2 and HaloTag to it. HaloTag-2xrGBD and 3xmTurquoise2-2xrGBD were created by digesting the backbone mNeonGreen-2xrGBD with AgeI and BsrGI and ligating the likewise digested inserts HaloTag and 3xmTurquoise2 to it. 1xrGBD-mNeonGreen-2xrGBD was created by PCR amplifying rGBD (primers: FW-5’ATATGCTAGCATGATCCTGGAGGACCTCAAT3’ and RV-5’ATATGCTAGCTCTAGAGCCTGTCTTCTCCAG3’), digesting it with NheI and ligating it to the likewise digested backbone mNeonGreen-2xrGBD. WASp(CRIB)-HaloTag-WASp(CRIB) was created by digesting the insert HaloTag with AgeI and BsrGI and ligating it to the likewise digested WASp(CRIB)-mScarlet-I-WASp(CRIB) backbone.

### Localized, constitutively active Rho GTPases

H2A-mTurquoise-Cdc42-G12V-ΔCaaX (addgene #176094), H2A-mTurquoise-Rac1-G12V-ΔCaaX (addgene #176095) and H2A-mTurquoise-RhoA-G14V-ΔCaaX (addgene #176097) plasmids were described previously (Mahlandt et al., 2021). mCherry-TOMM20-N1 was a gift from Michael Davidson (addgene #55146). To create TOMM20-mTurquoise-Cdc42-G12V-ΔCaaX, backbone H2A-mTurquoise2-Cdc42-G12V-ΔCaaX was digested with NdeI and BamHI and ligated to the likewise digested TOMM20. To create TOMM20-mTurquoise-Rac1-G12V-ΔCaaX, backbone H2A-mTurquoise-Rac1-G12V-ΔCaaX was digested with NdeI and BamHI and ligated to the likewise digested TOMM20. H2A-mTurquoise has been described previously (Goedhart et al., 2012)

### Rapamycin system

C1-Lck-FRB-mTuquoise2 was created by digesting C1-Lck-FRB-mTurquoise described before (Van Unen et al., 2015) and the insert mTurquoise2 (Goedhart et al., 2012) with AgeI and BsrGI and subsequently ligating the two fragments. Lck-FRB-ECFP(W66A) (addgene #67902) and YFP-FKBP12-p63RhoGEF(DH) have been described previously (Van Unen et al., 2015) as well as YFP-FKBP12-ITSN1(DHPH) (Reinhard et al., 2019). YFP-FKBP12-TIAM1(DHPH) was a gift from Tobias Meyer (addgene #20154).

### Other plasmids

The plasmid encoding histamine 1 receptor (H1R) was obtained from https://cdna.org/. The plasmids Lck-mTurquoise2-iLID (addgene #176125) and SspB-HaloTag-TIAM1(DHPH) (addgene # 176114) are described on addgene.

### Plasmid availability

The plasmids encoding dimericTomato-WASp(CRIB), also called wGBD, in a pLV backbone (addgene #176099) and C1 Backbone (addgene #191450), WASp(CRIB)-mTurquoise2-WASp(CRIB) (addgene #176136), WASp(CRIB)-HaloTag-WASp(CRIB) (addgene #191452), N-WASP(CRIB)-mScarlet-I-N-WASP(CRIB) (addgene #191451), TOMM20-mTurquoise-Cdc42-G12V-ΔCaaX (addgene #176121) and TOMM20-mTurquoise-Rac1-G12V-ΔCaaX (addgene #176124), mTurquoise2-3xrGBD in a C1 backbone (addgene #176100) and pEntr1a backbone (addgene 176137), HaloTag-3xrGBD (addgene #176108), HaloTag-2xrGBD (addgene #176109) are available on www.addgene.org.

### Cell Culture and Transfection

Henrietta Lacks (HeLa) cells (CCL-2, American Tissue Culture Collection) and Human Embryonic Kidney 293T (HEK293T) cells (CRL-3216, American Tissue Culture Collection) were cultured in Dulbecco’s modified Eagle’s medium + GlutaMAX™ (Giboc) with 10% fetal calf serum (Giboc) (DMEM + FCS) at 37°C and 7% CO_2_. The cell lines were routinely tested for mycoplasm contamination. For transfection 25 000 to 50 000 HeLa or Hek293T cells were seeded on round 24 mm ø coverslips (Menzel, Thermo Fisher Scientific) in a 6 well plate with 2 ml DMEM + FCS. The Transfection mix contained 1µl linear polyethylenimine (Polysciences) per 100 ng DNA, at a concentration of 1 mg/ml and a pH of 7.3, and 0.5 to 1 μg plasmid DNA per well, mixed with 200 μl OptiMEM (Thermo Fisher Scientific) per well. After 15 min incubation at room temperature the transfection mix was added to the cells, 24 h after seeding.

The SUM159 cells were a gift from Carol Otey (University of North Carolina at Chapel Hill, Chapel Hill, NC, USA). The SUM159 cells were cultured in Ham’s F12 (Cytiva) with 10% FBS, 5 μg/ml insulin (GIBCO), 1 μg/ml hydrocortisone (Sigma-Aldrich), and antibiotics. SUM159 were grown at 37°C and 5% CO2. Mycoplasma contamination was tested regularly by staining with Hoechst 33342 (AnaSpec Inc.). SUM159 cells were transfected with TransIT 2020 (Mirus).

### Imaging conditions

Cells were imaged between 24 to 48 h after transfection in an Attofluor cell chamber (Thermo Fischer Scientific), HeLa and Hek293T cell in 1 ml of Microscopy Medium (20 mM HEPES (pH=7.4),137 mM NaCl, 5.4 mM KCl, 1.8 mM CaCl2, 0.8 mM MgCl2 and 20 mM Glucose) at 37°C and BOECs in EGM+ at 37°C and 5% CO_2_. The HaloTag was stained, for at least 1 h prior to imaging, with a concentration of 150 nM of Janelia Fluor Dyes (JF) JF552 nm (red) or JF635 nm (far red) (Janelia Materials) in the culture medium. Rapamycin (LC Laboratories) diluted in DMSO was used at a final concentration of 100 nM, added to the cells during the imaging experiment and incubated for roughly 5 min. Human α-thrombin (HCT-0020, Haematologic technologies) diluted in PBS was used at a final concentration of 2 U/ml, directly added to the cells during the imaging. Phorbol ester Phorbol 12,13-dibutyrate (PDBu) (Sigma-Aldrich) was used at concentration of 1 µM SUM159 cells were imaged with 1:100 Oxyfluor reagent (Oxyrase Inc.) and 10 mM DL-lactate (Sigma-Aldrich) to reduce oxygen free radicals.

### Spinning disk microscopy

Spinning disk images were acquired with a Nikon Ti-E microscope equipped with a Yokogawa CSU X-1 spinning disk unit, a 60x objective (Plan Apo VC, oil, DIC, NA=1.4) and a 40x objective (Plan Fluor, oil, DIC, H/N2, NA=1.3), an Andor iXon 897 EMCCD camera, Perfect Focus System and the Nikon NIS elements software. CFPs were imaged using a 440 nm laser line, a triple dichroic mirror (440, 514, 561 nm) and a 460 – 500 nm emission filter. GFPs and YFPs were imaged using a 488 nm laser line, a triple dichroic mirror (405, 488, 561 nm) and a 500 nm long pass emission filter. RFPs were imaged using a 561 nm laser line, a triple dichroic mirror (405, 488, 561 nm) and a 600 – 660 nm emission filter.

### TIRFmicroscopy

TIRF microscopy images of HeLa and Hek 293T cells were acquired with a Nikon Ti-E microscope equipped with a motorized TIRF Illuminator unit, a 60× TIRF objective (60× Plan Apo, Oil DIC N2, NA=1.49, WD=120 μm), Perfect Focus System, with an Andor iXon 897 EMCCD camera and the Nikon NIS elements software. CFPs was imaged using the 440 nm laser line in combination with a tri split dichroic mirror (440, 488 and 561 nm). GFPs and YFPs were imaged using a 488 nm laser line in combination with a quad split dichroic mirror (405, 488, 561 and 640 nm) and a dual band pass emission filter (515–545 nm and 600–650 nm). RFPs were imaged using a 561 nm laser line in combination with a quad split dichroic mirror (405, 488, 561 and 640 nm) and a dual band pass emission filter (515–545 nm and 600–650 nm). Photo activation was achieved with a 440 nm laser line, intensity set to 20%, for 1 s repeated for every imaged frame.

TIRF microscopy images of SUM159 cells were acquired at a Nikon Eclipse Ti2 microscope, equipped with a Tokai Hit STX stage top incubator (set to 37°C and 5% CO2 for live imaging), Apo TIRF 60× NA 1.49 oil immersion objective, Hamamatsu ORCA-Flash 4.0 camera, Agilent laser unit (405-, 488-, 561-, and 647-nm), and NIS-Elements AR software.

### Confocal laser scanning microscopy

Confocal microscopy images were obtained at a Leica Sp8 (Leica Microsystems) equipped with a 63x objective (HC PL Apo, C2S, NA 1.40, oil) using unidirectional line scan at a scan speed of 400 Hz. Images were acquired with 1024x1024 pixel resolution and 16-bit color depth. The HaloTag stained with JF635 nm dye was imaged with the pinhole set to 1 Airy Unit, a 633 nm laser line and a HyD detector with an emission detection range of 647 nm to 710 nm and the gain set to 100 V.

### Analysis

#### Selection of candidates from GTPase interactor screen

Cdc42-GTP and Rac1-GTP binding proteins were selected from a mitchochondrial relocalization proximity biotinylation screen (Gillingham et al., 2019). From the 30 proteins with the highest WD score, for either binding Cdc42(G12V) or Rac1(Q61L), the ones that were only scoring for either Cdc42 or Rac1, were selected as sensor candidates. If available, plasmids containing the candidate’s sequences were obtained from addgene.

#### General image preparation

FIJI was used to analyze raw microscopy images (Schindelin et al., 2012). All images were background corrected. Example images were additionally adjusted for brightness and contrast and intensity was displayed as an inverted gray look up table, where darker colors represent higher intensity values.

Colocalization of mitochondrial tagged Rho GTPases and sensor candidates was judged by visual inspection shown in **Fig. 2A,B**.

To quantify nuclear colocalization of the sensor candidates with constitutively active nuclear localized Rho GTPases, an ROI of the nucleus was created by manually thresholding the intensity of H2A-mTurquoise2-Rho GTPase. The ROI representing the cytosol was created by enlarging the nucleus ROI by 0.5 µm and subtracting it by the nucleus ROI. The mean gray value was measured for the sensor candidates in the nucleus and cytosol ROI. The ratio of mean nuclear intensity to mean cytosolic intensity was calculated and plotted with the web tool ‘SuperPlotsofData’ shown in **Fig. 3B, 4B, 5G** (Goedhart, 2021).

The change in intensity was measured by either drawing an ROI for the cytosol or the plasma membrane by hand and measuring the mean gray value for the ROI. The ratio of the mean gray value pre stimulation to the mean gray value post stimulation was calculated and plotted as a percentage with the web tool ‘PlotsOfDifferences’ for **Fig. 5C,D,E, 6A, Sup 6C, Sup 7A,B** (Goedhart, 2019).

To measure the delta ratio of membrane to cytosol intensity, ROIs for the membrane and cytosol were drawn by hand and the mean gray value for these ROIs was measured. The ratio of pre over post stimulation for the ratio of the mean gray value for membrane over cytosol was calculated and plotted as a percentage with the web tool ‘PlotsofDifferences’ for **Fig. 5F** (Goedhart, 2019).

To measure the ratio of membrane over cytosol intensity over time, ROIs for the membrane and the cytosol were drawn by hand for each frame and the mean gray values for these ROIs were measured. The ratio of membrane over cytosol intensity was calculated, normalized by dividing each value by the value of the first frame and plotted with the web tool ‘PlotTwist’ for **Fig. 7B, Sup 7C** (Goedhart, 2020). In **Fig. 7D** solely the cytosolic intensity was plotted.

Images of **Fig. 7C** were flat field corrected, the background signal was subtracted, and bleaching was corrected with the ‘Exponential Fit Bleach Correction’ function in ImageJ.

#### Statistics

The 95% confidence interval was calculated by bootstrapping with the web tool ‘PlotsofData’ (Goedhart, 2021). The effect size that quantifies the difference and its distribution was calculated with the web tool ‘PlotsofDifferences’ (Goedhart, 2019).

## Acknowledgements

We want to thank Ronald Breedijk for the support at the van Leeuwenhoek Centre for Advanced Microscopy, Section Molecular Cytology, Swammerdam Institute for Life Sciences, University of Amsterdam.

## Author contribution

E.M. conceptualized, investigated, analyzed, visualized, and wrote the manuscript, G.K.-L. and R.G.-M. performed experiments, A.C. provided resources; J.G. acquired funding, supervised, conceptualized the project and reviewed and edited the manuscript.

## Data availability

The data generated during this study will be made available at Zenodo.org (after publication).

