## Supplementary figures and images for "Cell-based optimisation and characterisation of genetically encoded, location-based biosensors for Cdc42 or Rac activity"

### Movie 1

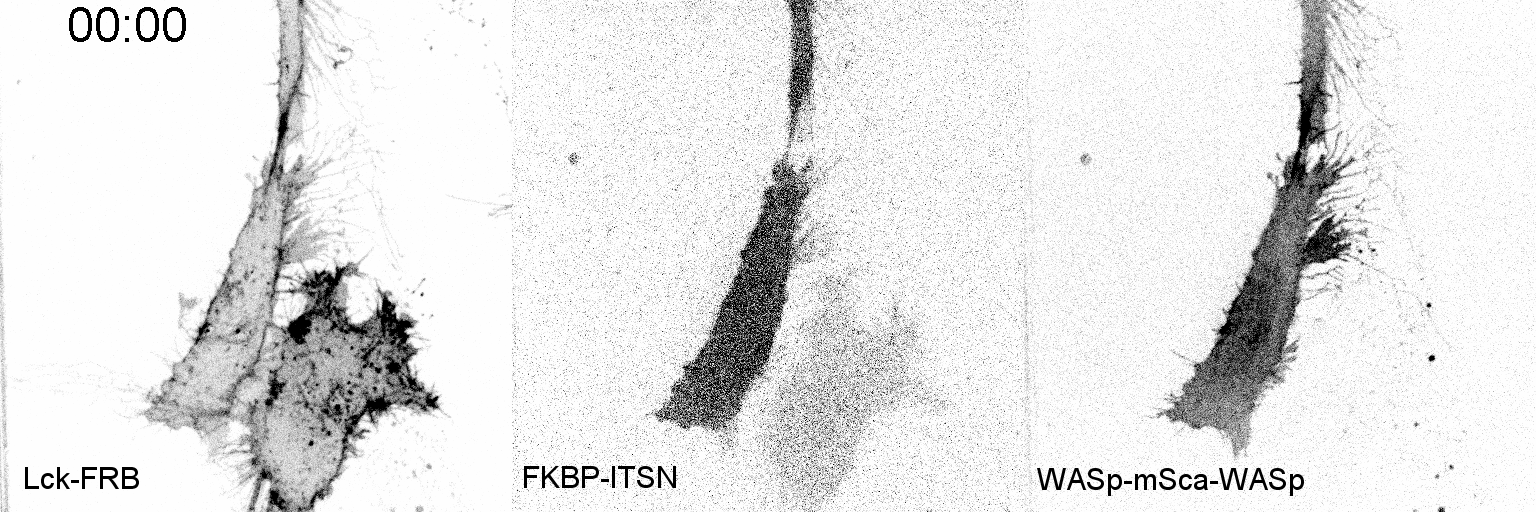

### Movie 2

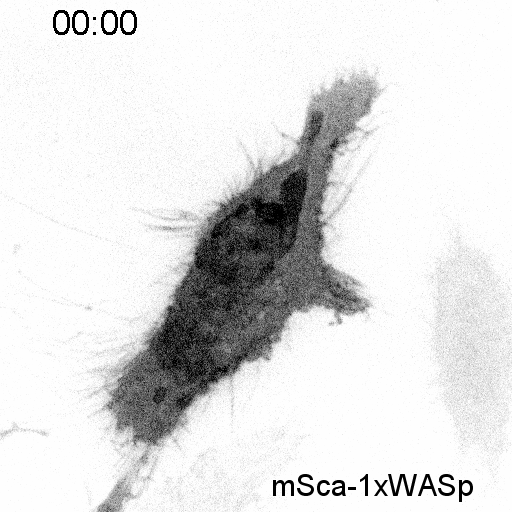

### Movie 3

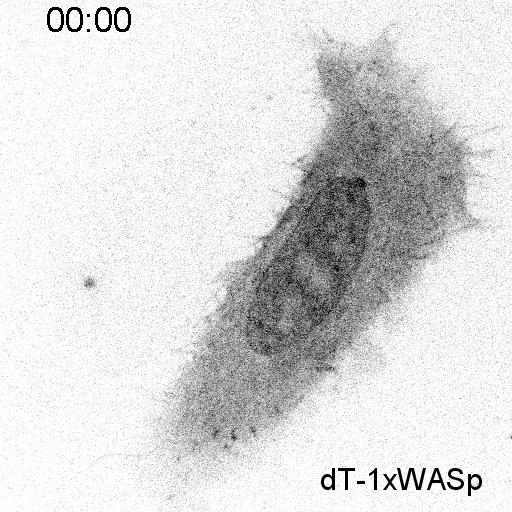

### Movie 4

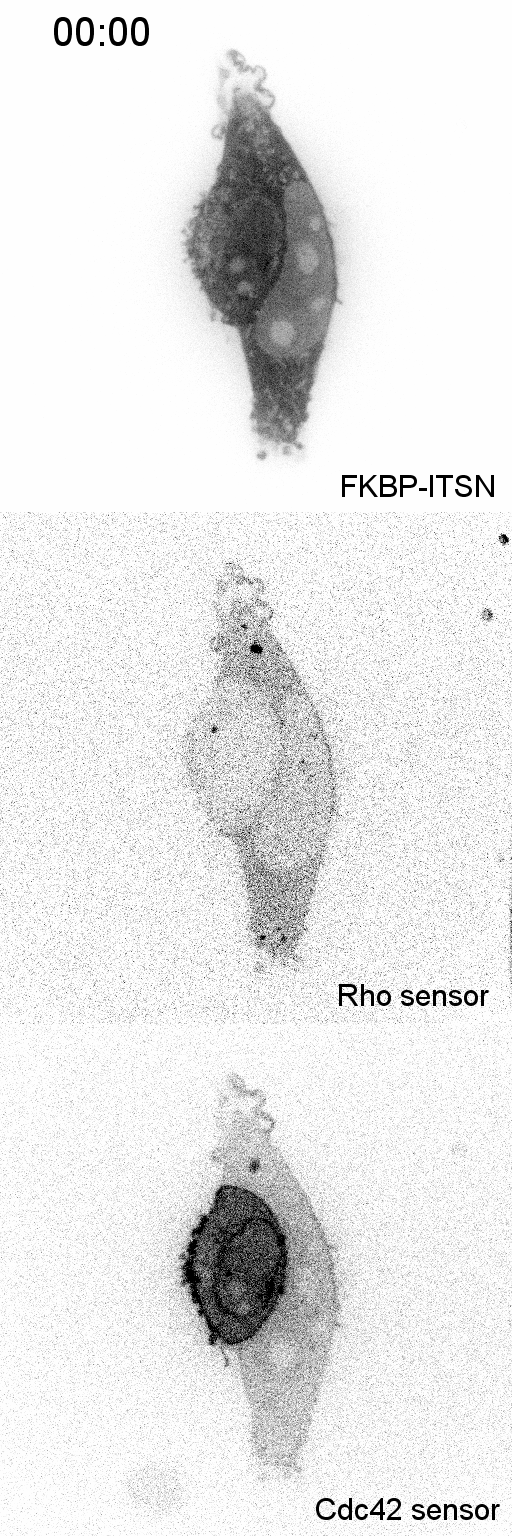

### Movie 5

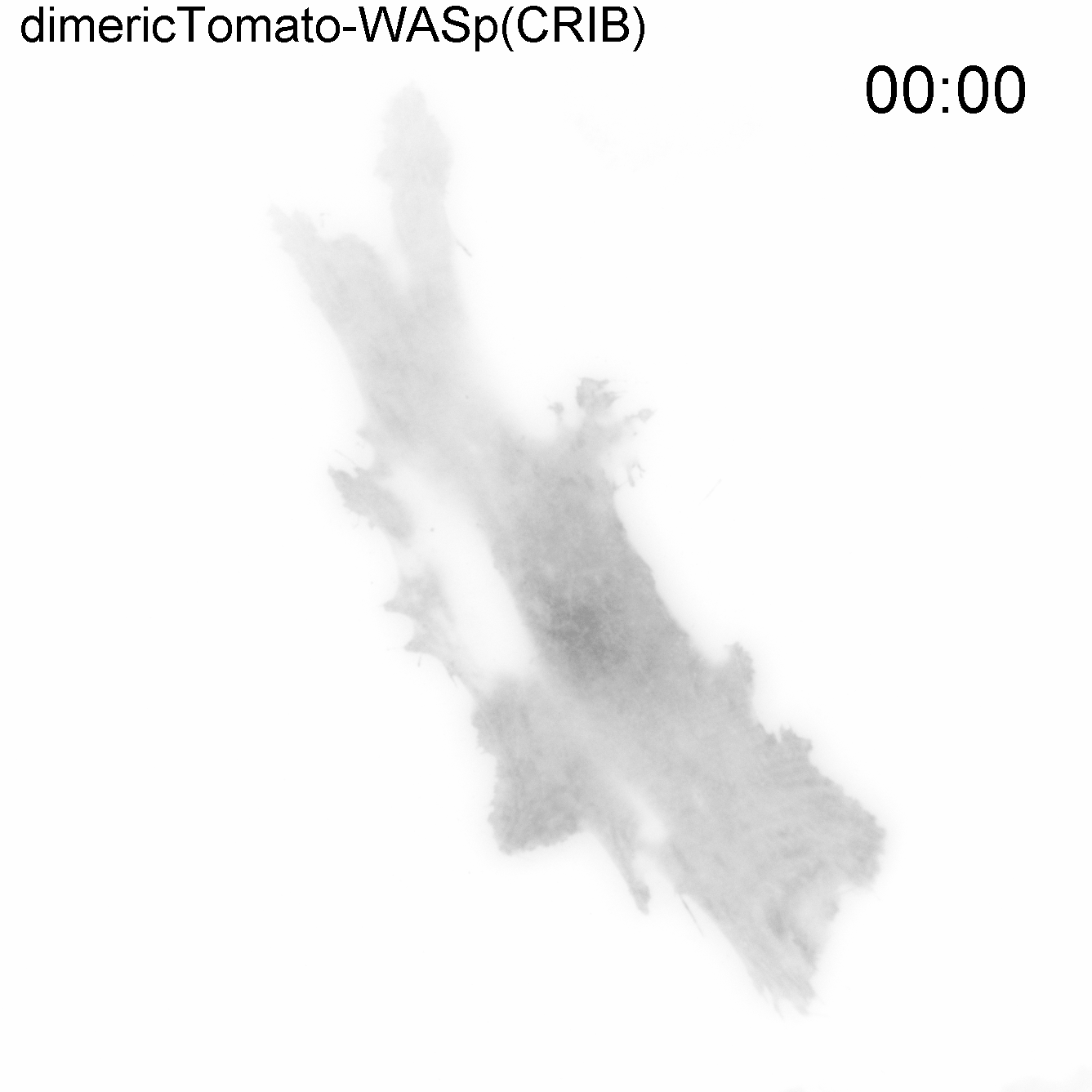

### Movie 6

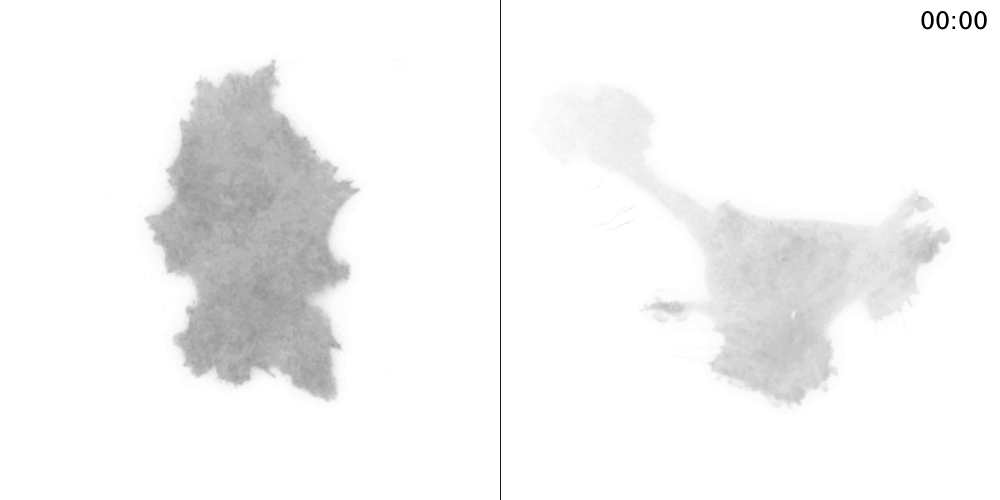

### Movie 7

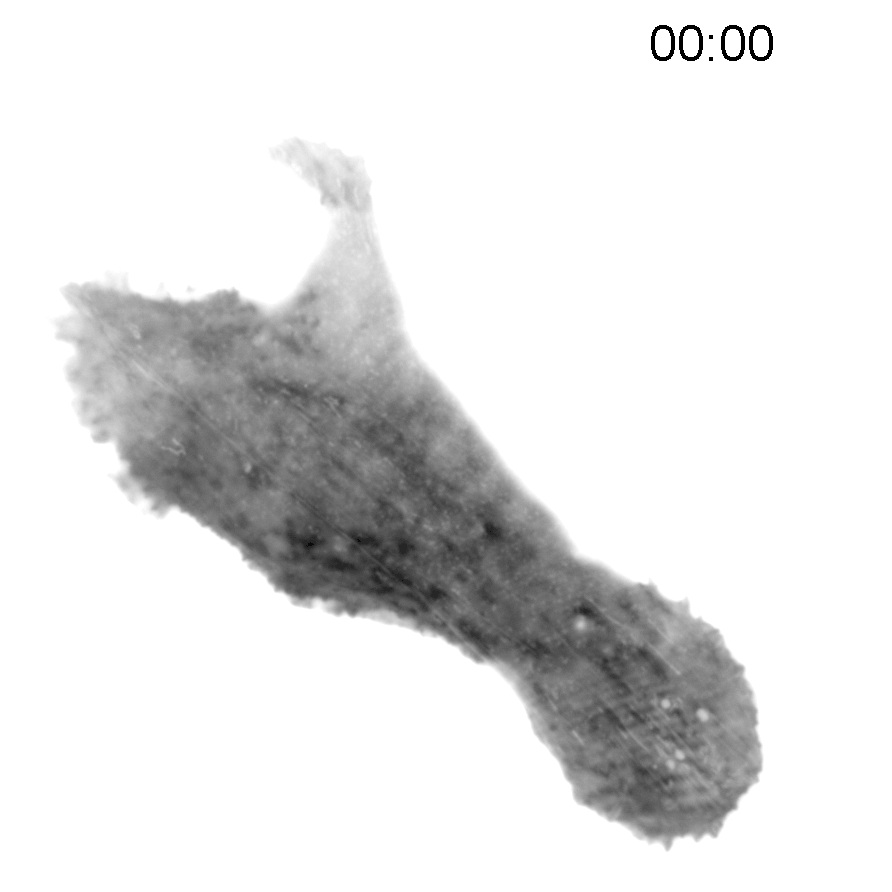
