## Supplemental data for "Cell-based optimisation and characterisation of genetically encoded, location-based biosensors for Cdc42 or Rac activity"

### Supplemental Figures

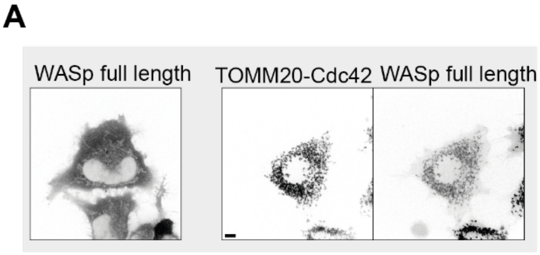


***Figure S1***. (***A***) *Spinning disk images of HeLa cells expressing mScarlet-I-WASp(full length) (left) and co-expressing of TOMM20-mTurquoise2-Cdc42(G12V)-ΔCaaX (middle) and mScarlet-I-WASp(full length) (right). Scale bar: 10 µm.*

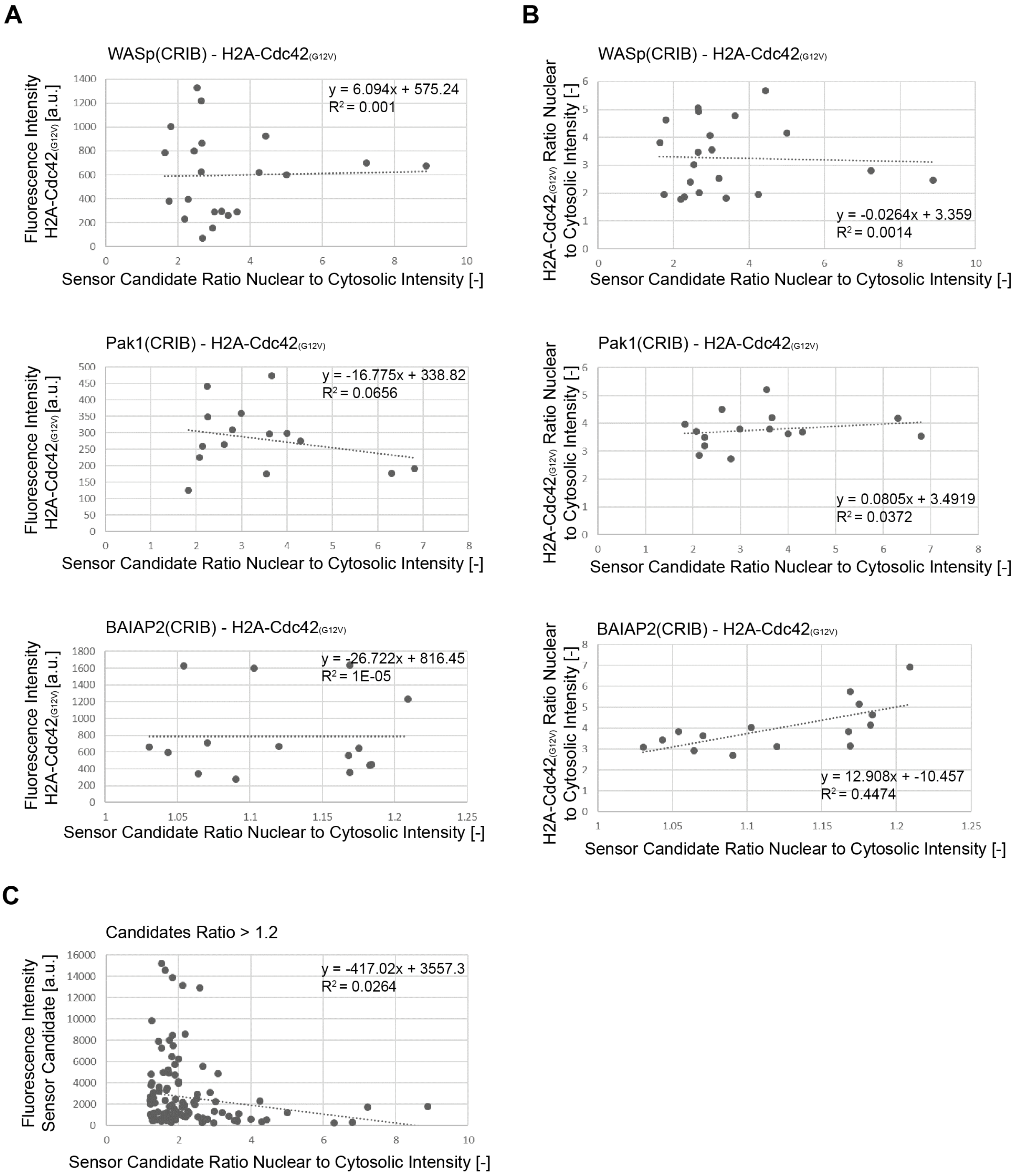


***Figure S2****. (****A****) Plots of the correlation between expression level of H2A-Cdc42(G12V)-ΔCaaX, represented by fluorescence intensity, and sensor candidate nuclear to cytosolic intensity, at the example of WASp(CRIB), Pak1(CRIB) and BAIAP2(CRIB), for the data shown in* ***Fig. 3B****. The general linear model (top, right corner) was fit to the data and is indicated as a dashed line. (****B****) Plots of the correlation between H2A-Cdc42(G12V)-ΔCaaX nuclear to cytosolic intensity and sensor candidate nuclear to cytosolic intensity, at the example of WASp(CRIB), Pak1(CRIB) and BAIAP2(CRIB), for the data shown in* ***Fig. 3B****. The general linear model (bottom, right corner) was fit to the data and is indicated as a dashed line. (****C****) Plots of the correlation between expression level of the sensor candidate, represented by fluorescence intensity, and sensor candidate nuclear to cytosolic intensity, for all candidates with a ratio above 1.2, for the data shown in* ***Fig. 3B****. The general linear model (top, right corner) was fit to the data and is indicated as a dashed line.*

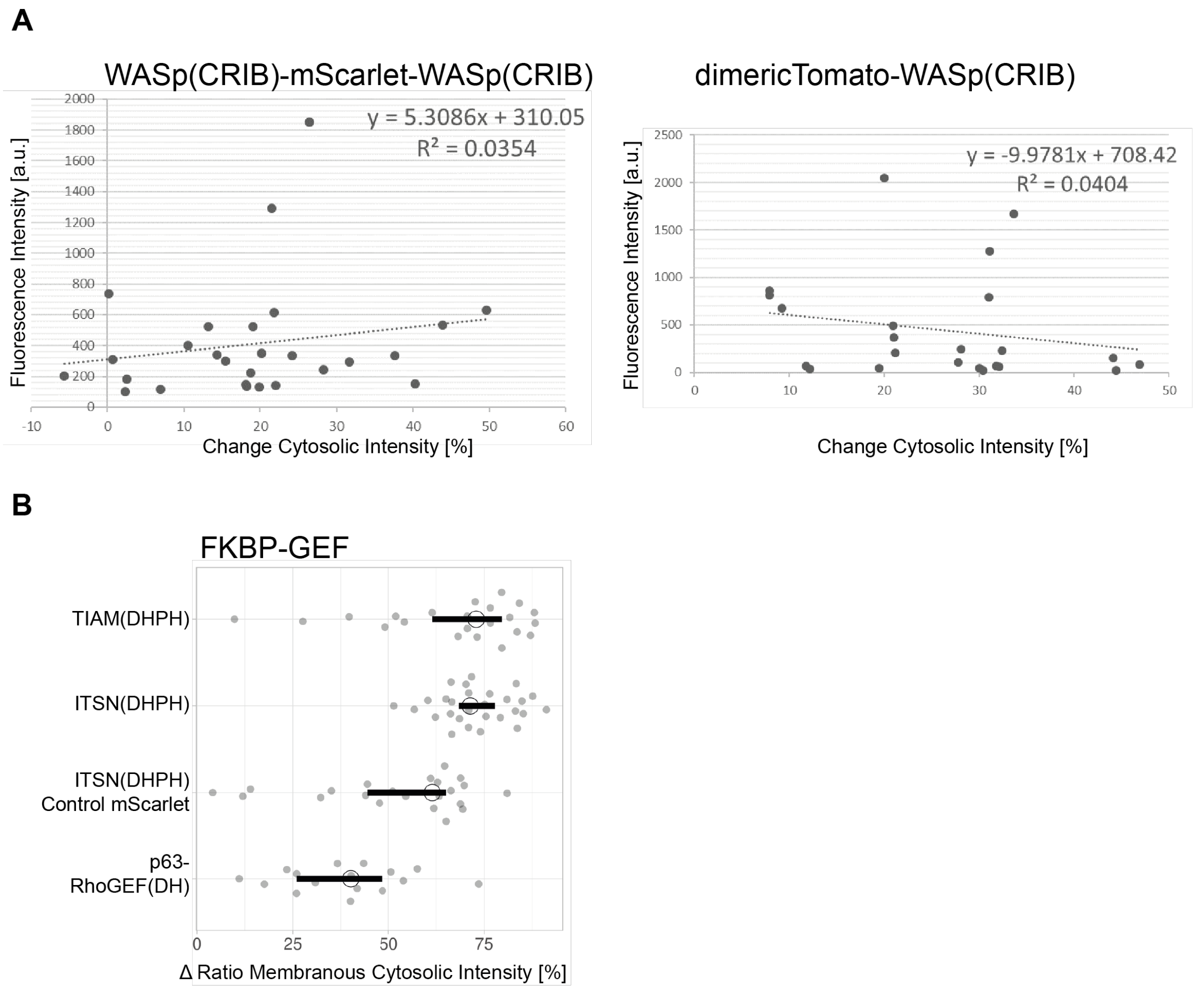


***Figure S3**. (**A**) Plots of the correlation between fluorescence intensity and change in cytosolic intensity for cell expressing WASp(CRIB)-mScarlet-WASp(CRIB) or dimericTomato-WASp(CRIB) for the data shown in **Fig. 5D,E**. The general linear model (top right corner) was fit to the data and is indicated as a dashed line. (**B**) Relocation of the FKBP-GEF fusion protein represented by the change in ratio of membranous over cytosolic intensity measured in HeLa cells expressing Lck-FRB-mTurquoise2, dimericTomato-WASp(CRIB) or mScarlet-I as a control and either YFP-FKBP-ITSN1(DHPH), YFP-FKBP-TIAM1(DHPH) or YFP-FKBP-RhoGEFp63(DH). FKBP-GEF recruitment was stimulated with 100 nM rapamycin. The median is represented as a circle, the 95% confidence interval as a black line and each dot represents the measurement of a single cell. The number of cells measured in two experiments based on independent transfections is: mScarlet-I YFP-FKBP-ITSN1(DHPH) control=22, YFP-FKBP-TIAM1(DHPH)=22, YFP-FKBP-RhoGEFp63(DH)=16, YFP-FKBP-ITSN1(DHPH)=28.*

### Supplement

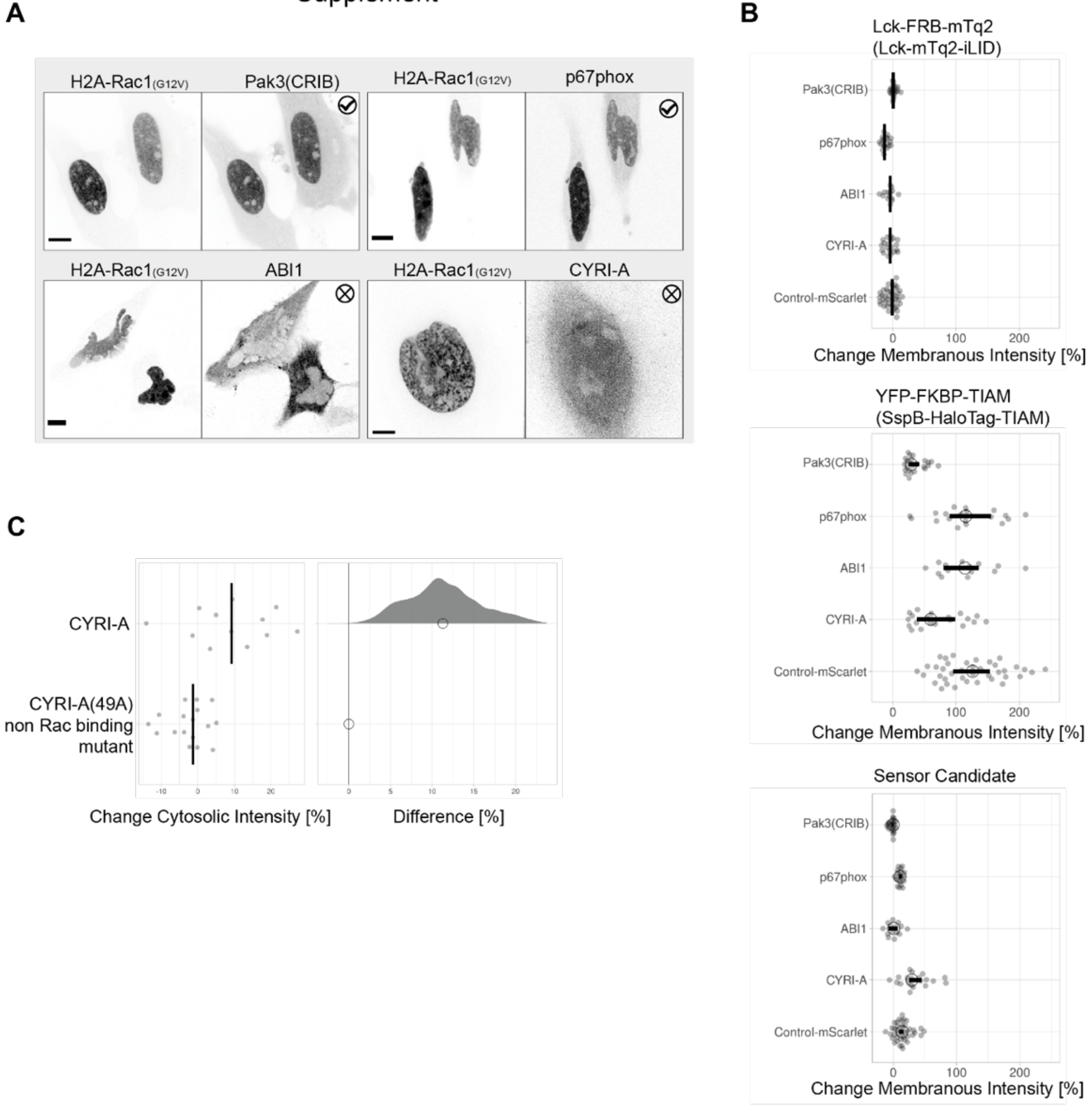


***Figure S4**. (**A**) Spinning disk images of HeLa cells expressing H2A-mTurquoise2-Rac1(G12V)-ΔCaaX (left) and either Rac sensor candidate mCherry-CYRI-A, mScarlet-I-ABI1 or p67phox-mCherry or EGFP-Pak3(CRIB) (right). Check mark indicating colocalization of candidate with nucleus by visual inspection. Scale bars: 10 µm. (**B**) Change in membranous intensity measured in HeLa cells expressing Lck-FRB-mTurquoise2, YFP-FKBP-TIAM1(DHPH) and either mCherry-CYRI-A, mScarlet-I-ABI1 or p67phox-mCherry, stimulated with 100 nM rapamycin. Additionally, HeLa cells expressing Lck-mTurquoise-iLID, SspB-HaloTag-TIAM1(DHPH) and EGFP-Pak3(CRIB), stimulated with 440 nm laser light at 1% intensity for 1 s in 20 s intervals. Measurements for all three channels, CFP, YFP and RFP. The median is represented as a circle and each dot represents the measurement of a single cell. The 95% confidence interval is represented by a bar. The number of cells measured in two experiments based on independent transfections is: ABI1=14, Control-mScarlet=41, CYRI-A=20, p67phox=19, Pak3(CRIB)=24. The same data is presented in **Fig. 6A**. (**C**) Relocation efficiency measured as a change in cytosolic intensity for CYRI-A and the non Rac binding mutant CYRI-A (49A) co-expressed in HeLa cells with Lck-FRB-mTurquoise2 and YFP-FKBP-TIAM1(DHPH) before and after the stimulation with 100 nM rapamycin. For the plot at the left, the median is represented as a black line and each dot represents the measurement of a single cell. The plot at the right shows the effect size relative to the non Rac binding mutant CYRI-A (49A). The bootstrap samples that are used to calculate the 95%CI of the effect size are shown as a distribution. The circle indicates the median. The number of cells measured is: CYRI-A=12, CYRI-A(49A)=17.*

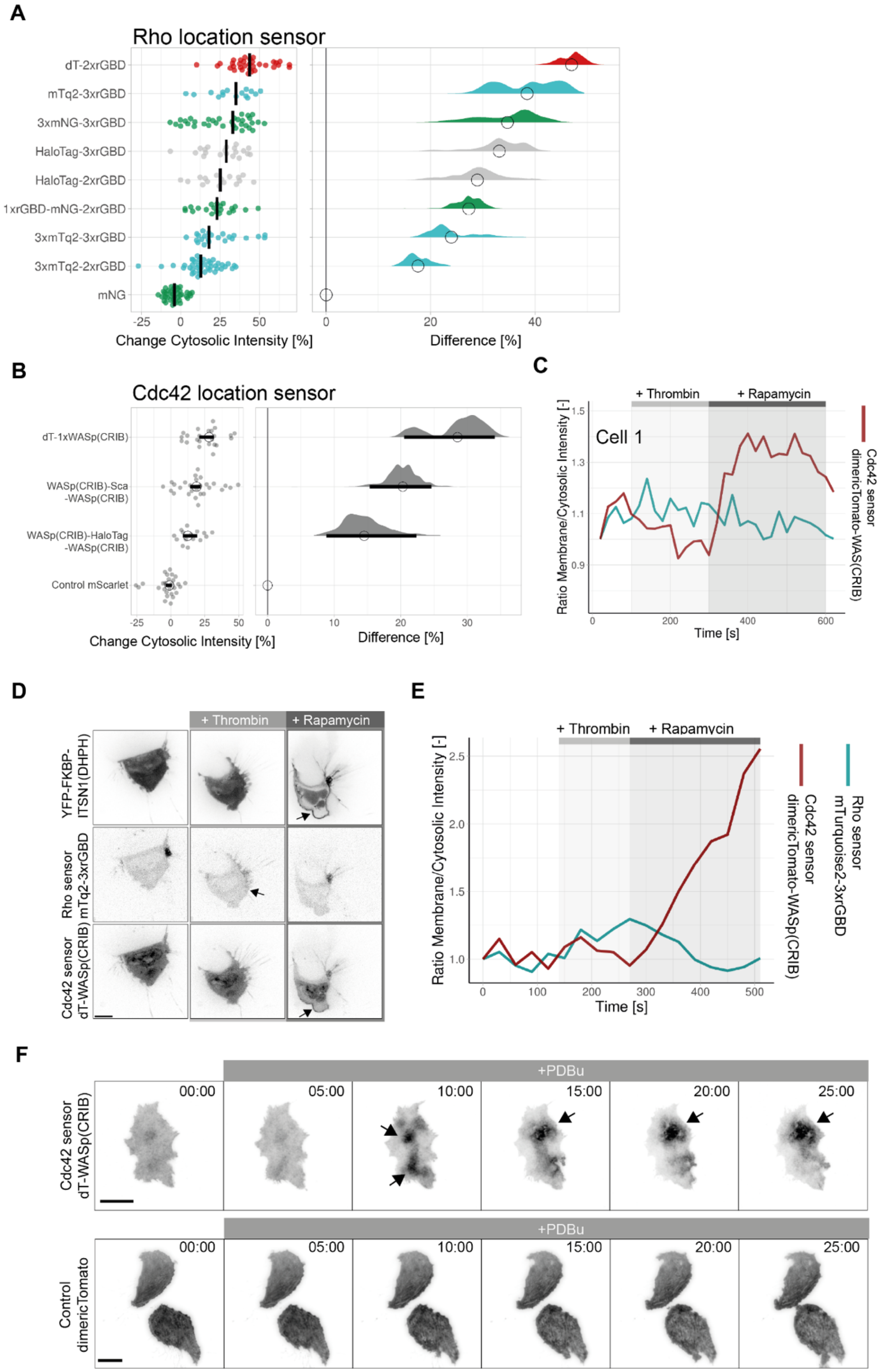

A
Rho location sensor
dT-2xrGBD
mTq2-3xrGBD
3xmNG-3xrGBD
HaloTag-3xrGBD
HaloTag-2xrGBD
1xrGBD-mNG-2xrGBD
3xmTq2-3xrGBD
3xmTq2-2xrGBD
mNG
Change Cytosolic Intensity [%]
Difference [%]
B
Cdc42 location sensor
dT-1xWASp(CRIB)
WASp(CRIB)-Sca -WASp(CRIB)
WASp(CRIB)-HaloTag -WASp(CRIB)
Control mScarlet
Change Cytosolic Intensity [%]
Difference [%]
C
+ Thrombin
+ Rapamycin
Cell 1
Ratio Membrane/Cytosolic Intensity [-]
Time [s]
Rho sensor mTurquoise2-3xrGBD
Cdc42 sensor dimericTomato-WAS(CRIB)
D
+ Thrombin
+ Rapamycin
YFP-FKBP-ITSN1(DHPH)
Rho sensor mTq2-3xrGBD
Cdc42 sensor dT-WASp(CRIB)
E
+ Thrombin
+ Rapamycin
Ratio Membrane/Cytosolic Intensity [-]
Time [s]
Rho sensor mTurquoise2-3xrGBD
Cdc42 sensor dimericTomato-WASp(CRIB)
F
+PDBu
00:00
05:00
10:00
15:00
20:00
25:00
Cdc42 sensor dT-WASp(CRIB)
Control dimericTomato

***Figure S5***. *(**A**) Relocation efficiency plotted as change in cytosolic intensity for CMV-mNeonGreen, CMVdel-3xmTurquoise2-2xrGBD, CMVdel-3xmTurquoise2-3xrGBD, CMVdel-1xrGBD-mNeonGreen-2xrGBD, CMVdel-HaloTag-2xrGBD, CMVdel-HaloTag-3xrGBD, 3xmNeonGreen-3xrGBD, CMVdel-mTurquoise2-3xrGBD and CMVdel-dimericTomato-2xrGBD in HeLa cells co-expressing the histamine 1 receptor (H1R), upon stimulation with 100 µM histamine. For the plot on the left, each dot represents an individual cell and the median of the data is shown as a black bar. The plot at the right shows the effect size relative to the mNeonGreen control. The bootstrap samples that are used to calculate the 95%CI of the effect size are shown as a distribution. The circle indicates the median. The number of cells measured in two experiments based on independent transfections is: 1xrGBD-mNG-2xrGBD=21, 3xmNG-3xrGBD=34, 3xmTq2-2xrGBD=42, 3xmTq2-3xrGBD=23, dT-2xrGBD=33, HaloTag-2xrGBD=16, HaloTag-3xrGBD=20, mNG=39, mTq2-3xrGBD=15. Data for CMV-mNeonGreen and CMVdel-dimericTomato-2xrGBD was taken from* (Mahlandt et al., 2021). *(**B**) Relocation efficiency plotted as change in cytosolic intensity for mScarlet-I, WASp(CRIB)-HaloTag-WASp(CRIB), WASp(CRIB)-mScarlet-I-WASp(CRIB) and dimericTomato-WASp(CRIB) in HeLa cells co-expressed with Lck-FRB-mTurquoise2 and YFP-FKBP-ITSN1(DHPH), stimulated with 100 nM rapamycin. For the plot on the left, each dot represents an individual cell and the median of the data is shown as a black bar. The plot at the right shows the effect size relative to the mNeonGreen control. The bootstrap samples that are used to calculate the 95%CI of the effect size are shown as a distribution. The circle indicates the median. The number of cells measured is: mScarlet-I=27, WASp(CRIB)-HaloTag-WASp(CRIB)=15, WASp(CRIB)-mScarlet-I-WASp(CRIB)=27, dimericTomato-WASp(CRIB)=23. Data for mScarlet-I, WASp(CRIB)-mScarlet-I-WASp(CRIB) and dimericTomato-WASp(CRIB) is also shown in **Fig. 6C,D,E**. (**C**) Time trace of the ratio membrane over cytosolic intensity for the in **Fig. 7B** depicted cell 1 for the Rho sensor mTurquoise2-3xrGBD (blue) and the Cdc42 sensor dimericTomato-WASp(CRIB). Thrombin addition is indicated by a light grey bar. Rapamycin addition is indicated by a dark grey bar. **D**) Spinning disk images of Hek 293T cells expressing Lck-FRB-ECFP(W66A)-black mutant, YFP-FKBP-ITSN1(DHPH) (top), Rho sensor mTurquoise2-3xrGBD (middle) and the Cdc42 sensor dimericTomato-WASp(CRIB) (bottom), first stimulated with 2 U/ml human α-thrombin, and then with 100 nM rapamycin. Arrows indicate intensity increase at the plasma membrane. Scale bar: 10 µm. (**E**) Time trace of the ratio membrane over cytosolic intensity for the in **D** depicted cell for the Rho sensor mTurquoise2-3xrGBD (blue) and the Cdc42 sensor dimericTomato-WASp(CRIB). Thrombin addition is indicated by a light grey bar. Rapamycin addition is indicated by a dark grey bar. **F**) TIRF microscopy images of a SUM159 cell expressing the Cdc42 sensor dimericTomato-WASp(CRIB) (upper panel) or dimericTomato as a control (lower panel), stimulated with PDBu (1 µM), as indicated by the grey bar. Arrows indicate the local signal accumulation of the Cdc42 sensor. Time is given in min:s from the beginning of the recording. Scale bar: 25 µm.*

#### Movie Legends:

**Movie 1. The WASp(CRIB)-mScarlet-I-WASp(CRIB) Cdc42 sensor relocating in a HeLa cell**. Spinning disk time lapse movie of a HeLa cell co-expressing Lck-FRB-mTurquoise2 (left), YFP-FKBP-ITSN1(DHPH) (middle) and WASp(CRIB)-mScarlet-I-WASp(CRIB), stimulation with 100 nM rapamycin after 40 s. The time stamper represents min:s from the beginning of the recording.

**Movie 2. The mScarlet-I-1xWASp(CRIB) Cdc42 sensor relocating in a HeLa cell**. Spinning disk microscopy images of a HeLa cell co-expressing Lck-FRB-mTurquoise2 (not shown), YFP-FKBP-ITSN1(DHPH) (not shown) and mScarlet-I-1xWASp(CRIB), stimulated with 100 nM rapamycin at 1:40 min. The time stamper represents min:s from the beginning of the recording.

**Movie 3. The dimericTomato-1xWASp(CRIB) Cdc42 sensor relocation in a HeLa cell**. Spinning disk microscopy images of a HeLa cell (not the cell depicted in Fig. 5B) co-expressing Lck-FRB-mTurquoise2 (not shown), YFP-FKBP-ITSN1(DHPH) (not shown) and dimericTomato-1xWASp(CRIB), stimulated with 100 nM rapamycin at 1:20 min. The time stamper represents min:s from the beginning of the recording.

**Movie 4. Co-imaging of the optimized Cdc42 and Rho relocation sensors.** Spinning disk images of Hek 293T cells co-expressing Lck-FRB-ECFP(W66A)-black mutant (not shown), YFP-FKBP-ITSN1(DHPH) (top), Rho sensor mTurquoise2-3xrGBD (middle) and the Cdc42 sensor dimericTomato-WASp(CRIB) (bottom), first stimulated with 2 U/ml human α-thrombin, and then with 100 nM rapamycin. The time stamper represents min:s from the beginning of the recording.

**Movie 5. Visualization of local Cdc42 activity**. TIRF microscopy time-lapse of a SUM159 cell expressing the Cdc42 sensor dimericTomato-WASp(CRIB) stimulated with PDBu as indicated. Time is given in min:s from the beginning of the recording.

**Movie 6. Visualization of local Cdc42 activity**. A composite of two TIRF microscopy time-lapses of a SUM159 cell expressing the Cdc42 sensor dimericTomato-WASp(CRIB) stimulated with PDBu as indicated. Time is given in min:s from the beginning of the recording.

**Movie 7. Control for the visualization of local Cdc42 activity.** TIRF microscopy time-lapse of a SUM159 cell expressing dimericTomato as a control stimulated with PDBu as indicated. Time is given in min:s from the beginning of the recording.
